# Nanoscale spatial-omics via contrastive embedding of single-molecule localisation data

**DOI:** 10.1101/2025.10.07.679170

**Authors:** S. Shirgill, L.G. Jensen, M.C. Wales, A. Kaminer, K. Savoye, R. Peters, M. Heilemann, R. Henriques, S.F. Lee, P. Rubin-Delanchy, D.J. Nieves, D.M. Owen

**Affiliations:** Department of Immunology and Immunotherapy, School of Infection, Inflammation and Immunology, College of Medicine and Health, University of Birmingham, UK; School of Mathematics, College of Engineering and Physical Sciences, University of Birmingham, UK; Department of Mathematics, Aarhus University, Denmark; Institute of Physical and Theoretical Chemistry, Goethe-University Frankfurt, Germany; International Max Planck Research School (IMPRS) on Cellular Biophysics, Frankfurt am Main, Germany; School of Physics and Astronomy, College of Engineering and Physical Sciences, University of Birmingham, UK; School of Mathematical and Physical Sciences, University of Sheffield, UK; Instituto de Tecnologia Química e Biológica António Xavier, Universidade NOVA de Lisboa, Portugal; Yusuf Hamied Department of Chemistry, University of Cambridge, UK; School of Mathematics, University of Edinburgh, UK; Centre of Membrane Proteins and Receptors, University of Birmingham, UK

## Abstract

Omics approaches have transformed biology by characterising cells at the genomic, transcriptomic and proteomic levels. An additional layer of cellular information is provided by the nanoscale spatial organisation of molecules, which can be measured using super-resolution microscopy. We present a self-supervised contrastive learning framework that embeds single-molecule localisation microscopy (SMLM) point patterns into a learned representation of nanoscale organisation. Across simulated and experimental data, the representation distinguishes diverse spatial architectures, captures graded changes in organisation and supports unsupervised comparison of experimental conditions. The framework distinguishes proteins with different nanoscale distributions and captures variation associated with cell type and pharmacological perturbation. In a direct comparison with six conventional DBSCAN- and Ripley-derived spatial descriptors, the learned representation showed stronger correspondence with protein identity during unsupervised clustering, while conventional descriptors retained substantial discriminatory information. Together, these findings establish contrastive representation learning as a complementary, scalable framework for comparative nanoscale phenotyping and hypothesis generation.

## Introduction

Omics approaches enable comprehensive characterisation of biological systems at the molecular level. One of the most well-established is genomics, where advances in sequencing technologies and the availability of large public reference datasets have enabled systematic studies of genetic variation and genome-wide association with complex traits^1,2^. Single-cell transcriptomics has provided insights into developmental biology and the discovery of new cell types and states^3,4^, while advances in computational protein structure prediction, exemplified by AlphaFold, have transformed structural biology^5^.

Beyond molecular composition, spatial context provides an additional dimension of cellular characterisation. Spatial transcriptomics and multiplexed tissue imaging have demonstrated the value of mapping RNA and protein distributions within their native cellular environments^6,7^. Here, we focus on a complementary level of spatial information: the nanoscale arrangement of single-molecule localisations within subcellular regions, rather than the spatial distribution of molecular abundances across cells and tissues. Understanding this organisation is a central motivation behind fluorescence microscopy, one of the most widely used research techniques in cell biology. However, conventional diffraction-limited fluorescence microscopy typically cannot resolve structures separated by less than approximately 200 nm laterally^8^. Developments in fluorescence nanoscopy, including single-molecule localisation microscopy (SMLM) and MINFLUX, alongside complementary ultrastructural approaches such as gold-labelled electron microscopy, have enabled molecular organisation to be investigated at substantially smaller length scales^8–10^. Within this landscape, SMLM is particularly well suited to the quantitative investigation of protein organisation at the nanoscale^8,11^.

SMLM analysis encompasses a range of spatial-statistical and computational approaches, including Ripley’s K-function and pair-correlation analysis^12,13^, clustering and segmentation methods such as DBSCAN and Voronoi tessellation^13–15^, and machine-learning approaches for cluster detection and morphological characterisation^16–18^. These methods are powerful and provide biologically interpretable measurements, but typically require decisions about which spatial features, parameters or training targets to investigate. This can complicate exploratory comparisons across heterogeneous experimental datasets. Here, we apply self-supervised contrastive learning to derive a shared representation of SMLM point patterns without requiring biological class labels or the manual preselection of spatial descriptors^19^.

We use contrastive learning to map SMLM datasets into a high-dimensional representation in which organisationally similar point patterns tend to occupy nearby regions, enabling systematic comparison at scale. Similar approaches are used in single-cell transcriptomics, where high-dimensional gene expression profiles are represented in feature space and subsequently explored using dimensionality reduction methods such as UMAP^20,21^. Importantly, we do not assume that distance within the learned representation is linearly proportional to a specific physical or biological quantity. Instead, the high-dimensional representation provides a common feature space for downstream analyses including clustering, classification and similarity measurement, while UMAP is used only for qualitative two-dimensional visualisation. We therefore refer to this framework as nanoscale spatial-omics: a scalable approach for comparative analysis of nanoscale protein organisation across large SMLM datasets^22^.

## Results

To study nanoscale protein organisation, SMLM datasets were divided into standardised 3×3 μm regions of interest (ROIs) for input into the trained contrastive encoder. Each ROI was converted into a fixed-resolution (120×120 bins/25 nm per bin) spatial representation. ROIs containing related spatial organisations tend to occupy similar regions of this learned representation, enabling downstream analyses such as clustering, classification and calculation of representation-space similarity. Unless otherwise stated, K-means clustering was performed directly in the full 128-dimensional representation (128D). For visualisation only, representations were projected into two dimensions using UMAP^21^ with fixed random seeds. Consequently, apparent distances or boundaries in the two-dimensional UMAP projections were not used for quantitative clustering or interpreted as direct measures of biological difference.

To assess the relationship between the synthetic training distribution and experimental SMLM data, we jointly embedded Perlin training patterns and the five experimental protein datasets (**Supplementary Figure S1**). The experimental representations showed substantial overlap with the synthetic training distribution, although coverage varied between protein architectures. We quantified this using a k-nearest-neighbour support criterion defined from distances between training samples in the full 128D representation. Membrane-associated proteins were predominantly contained within the training-supported regions, whereas microtubule and especially actin ROIs extended beyond them. These findings indicate that Perlin-based training provides substantial, but not exhaustive, coverage of the experimental representation space, while the encoder also captures structural organisation in experimental patterns that are less densely represented in the training distribution.

The encoder was trained using an adapted SimCLR-style contrastive framework^19^ (**Figure 1b**). Training point patterns were generated procedurally using Perlin noise^23,24^, providing a broad distribution of heterogeneous spatial organisations without assigning biological class labels. Perlin noise was used as a flexible training distribution rather than as a model of any particular protein architecture; the purpose was to expose the encoder to variation in spatial scale, density and morphology before application to experimental data. Two independently generated point patterns sharing the same generative parameters were treated as a positive pair. Each member of the pair was passed through the shared encoder and a projection head used only during training, and the contrastive objective encouraged related representations to agree while separating patterns generated from different parameter settings. After training, the projection head was discarded and the encoder alone was used to generate 128D representations of simulated and experimental ROIs.

**Figure 1:**
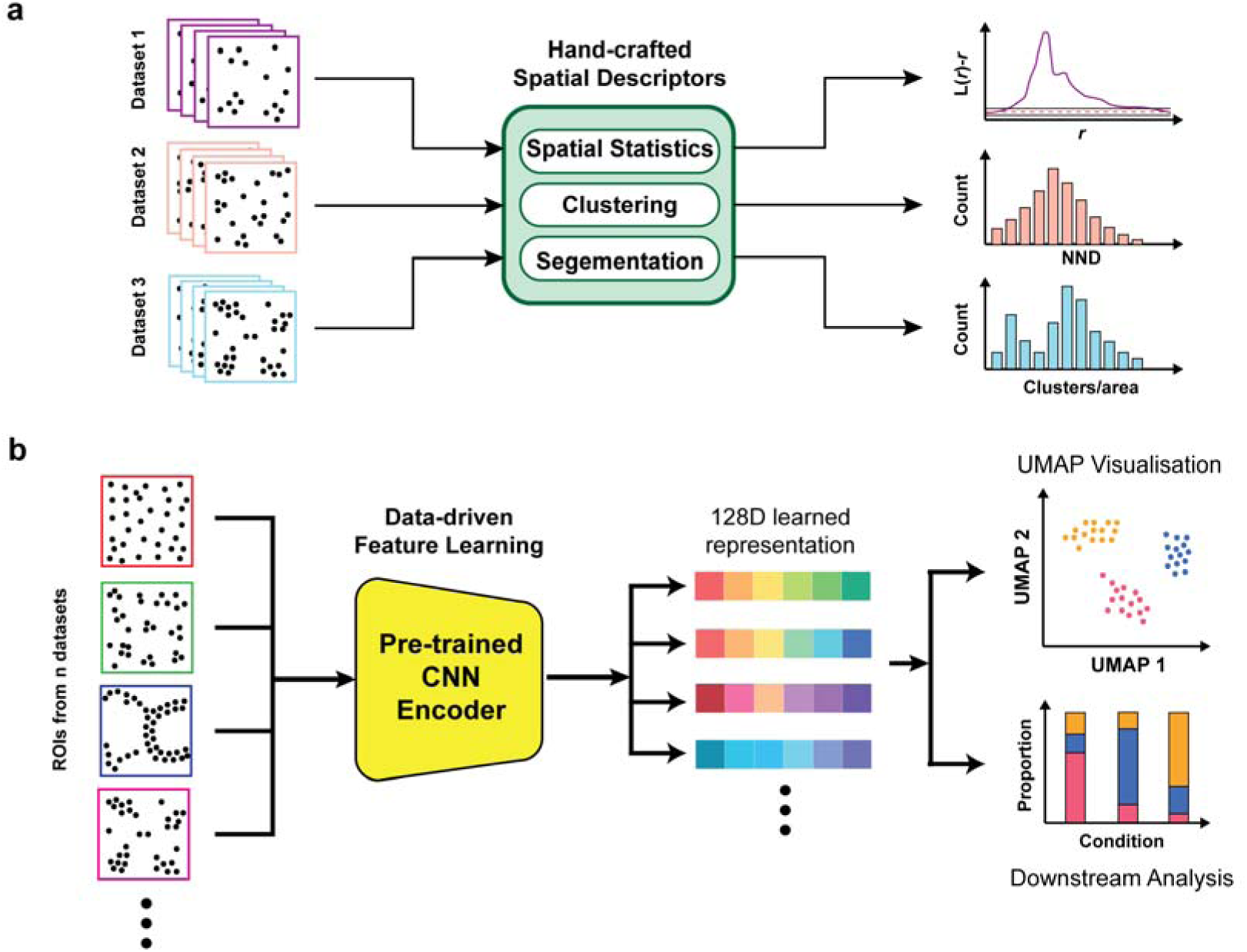
Overview of conventional and contrastive-learning approaches for SMLM point-pattern analysis. **a)** Conventional analysis of single-molecule localisation microscopy (SMLM) data relies on predefined spatial descriptors to characterise molecular organisation. Approaches such as spatial statistics, clustering and segmentation quantify selected properties of localisation patterns, producing measurements including Ripley-derived functions, nearest-neighbour distance distributions and cluster densities. These descriptors provide interpretable measures of spatial organisation, but require the analyst to select the spatial features and parameters of interest in advance. **b)** In our proposed contrastive-learning framework, a convolutional neural network (CNN), pretrained on procedurally generated Perlin-noise patterns, maps experimental SMLM regions of interest (ROIs) into a shared 128-dimensional (128D) learned representation without requiring experimental condition labels during training. The resulting embeddings can be visualised using uniform manifold approximation and projection (UMAP) and subjected to downstream analyses, including clustering of the 128D representations and comparisons of cluster composition across experimental conditions.

To evaluate whether the learned representation captured controlled differences in nanoscale spatial organisation, we first analysed simulated 3×3 µm ROIs containing different numbers of randomly positioned Gaussian clusters (**Figure 2** and **Table 1**). ROIs containing 2, 5, 9, 10 or 20 clusters were encoded into the 128D representation and projected using UMAP for visualisation. Most simulated conditions occupied distinct regions of the projection, while the 9- and 10-cluster conditions showed greater overlap, consistent with their comparatively similar spatial organisation. K-means clustering performed directly in the 128D representation with k = 5 similarly produced clusters enriched for individual simulation conditions (ARI = 0.807), with the greatest mixing occurring between closely related parameter values.

**Figure 2:**
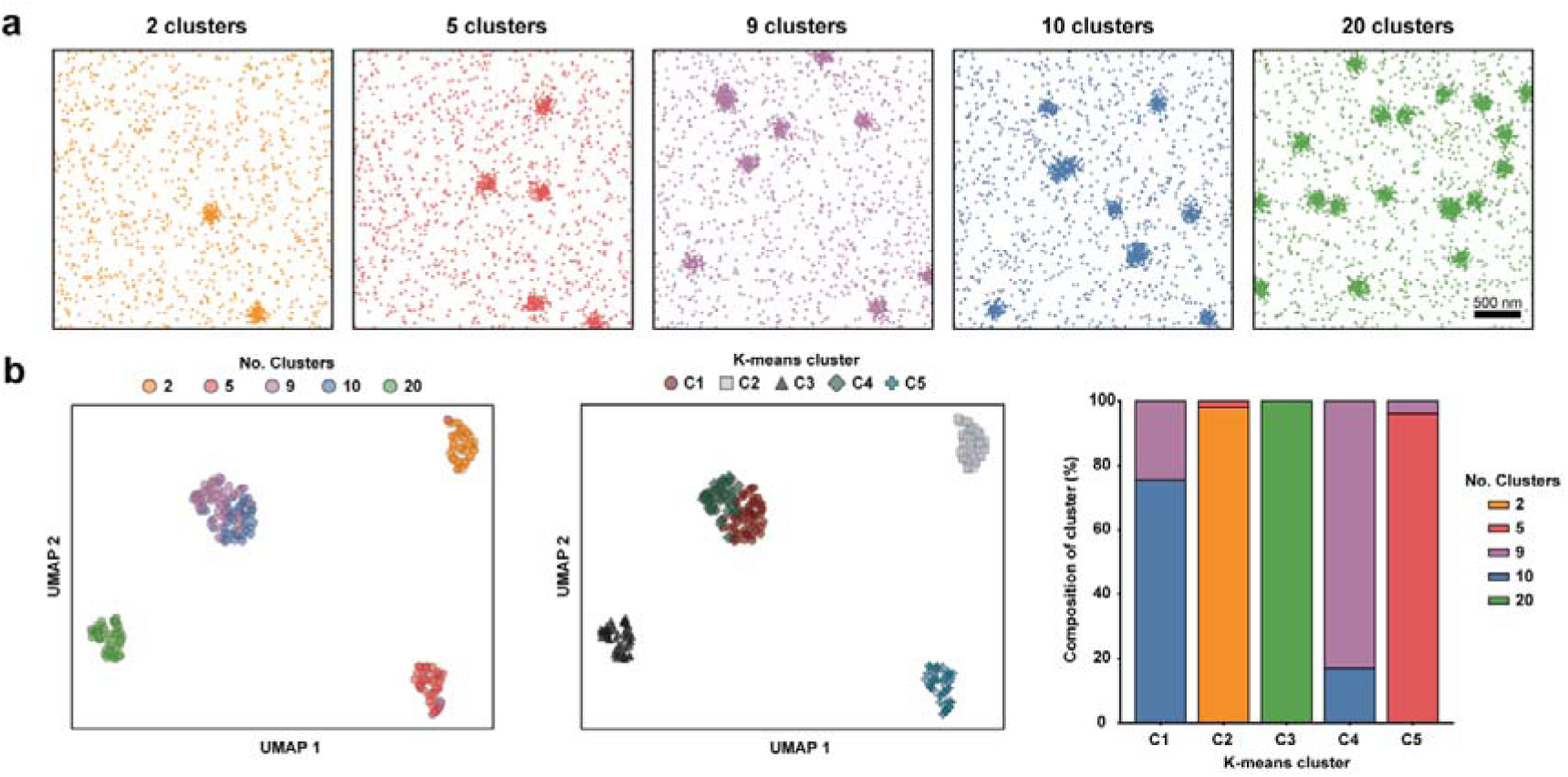
Contrastive learning discriminates simulated point patterns according to their nanoscale organisation. **a)** Representative 3×3 µm simulated ROIs containing 2, 5, 9, 10 or 20 randomly positioned Gaussian clusters. Scale bar, 500 nm. **b)** ROIs were encoded into the 128D learned representation and visualised using UMAP, pseudo-coloured either by simulation condition (left) or by K-means cluster assignment (centre). K-means clustering was performed in the full 128D representation with k = 5, matching the number of simulated conditions. The percentage condition composition of each K-means cluster is shown (right). Each simulated condition comprised 50 independently generated ROIs (n = 250 total).

**Table 1:** Parameters used in the independent simulation experiments.

| Simulation experiment | Parameter varied | Values tested | Figure |
|---|---|---|---|
| **Gaussian-cluster number** | Number of clusters per ROI | 2, 5, 9, 10, 20 | **Fig. 2** |
| **Cluster localisation density** | Localisations per Gaussian cluster | 10, 50, 90, 100, 200 | **Fig. S2b** |
| **Gaussian-cluster width** | Isotropic Gaussian standard deviation (nm) | 30, 35, 40, 49, 50, 100, 200 | **Fig. S2c** |
| **Background variation** | Simulation background-density parameter | 0.1, 0.4, 0.5, 0.9 | **Fig. S2d** |
| **Relative fibre density** | Fibre-to-cluster localisation-count ratio | 1, 2, 3, 5, 10, 20× | **Fig. S2a** |
| **Extended fibre-density sweep** | Fibre-to-cluster localisation-count ratio | 1, 2, 3, 5, 10, 20, 50, 100, 200, 500× | **Fig. S3** |

We next extended this analysis across several independent simulation families in which relative fibre density, localisation number per cluster, Gaussian-cluster standard deviation or background density were systematically varied (**Supplementary Figure S2**). Each perturbation generated an ordered response within the learned representation, while joint embedding of all simulation families showed that different classes of spatial organisation occupied distinct but related regions of a common representation (**Supplementary Figure S2e**). These results indicate that the encoder captures multiple aspects of point-pattern organisation rather than responding exclusively to a single parameter such as localisation density.

To characterise the extent to which density within structures can saturate separation in the learned representation, we extended the simulations in **Supplementary Figure S2a** to sweep from 1x to 500x relative fibre density (**Supplementary Figure S3**). Median cosine distance from the 1x reference increased strongly between 1x and approximately 10–20x fibre density, but changed little thereafter, remaining approximately constant between 50x and 500x. Thus, the learned representation retains sensitivity to low- and intermediate-range changes in fibre density but approaches a saturated regime for extremely fibre-dominated point patterns. This behaviour defines an empirical limit of sensitivity rather than implying that the underlying simulated structures themselves become identical.

Next, we evaluated whether the learned representation could distinguish experimental SMLM datasets corresponding to different proteins (**Figure 3**). Datasets for microtubules, actin, KIR2DL1, TIGIT and Lck were obtained from the nano-org database^22^ or acquired inhouse and comprised a range of spatial organisations, cell types and labelling conditions. Representative full fields of view and the corresponding 3×3 µm ROIs illustrate the substantial structural diversity present across these datasets (**Figure 3a**). In the learned representation, the five proteins occupied distinct but partially overlapping regions of the UMAP visualisation (**Figure 3b**). K-means clustering performed directly in the 128D representation with k = 5 produced moderate agreement with protein identity (ARI = 0.532, 95% CI 0.497–0.573; permutation p = 0.0002), indicating that protein-specific nanoscale organisation is captured by the learned representation despite substantial within-condition heterogeneity.

**Figure 3:**
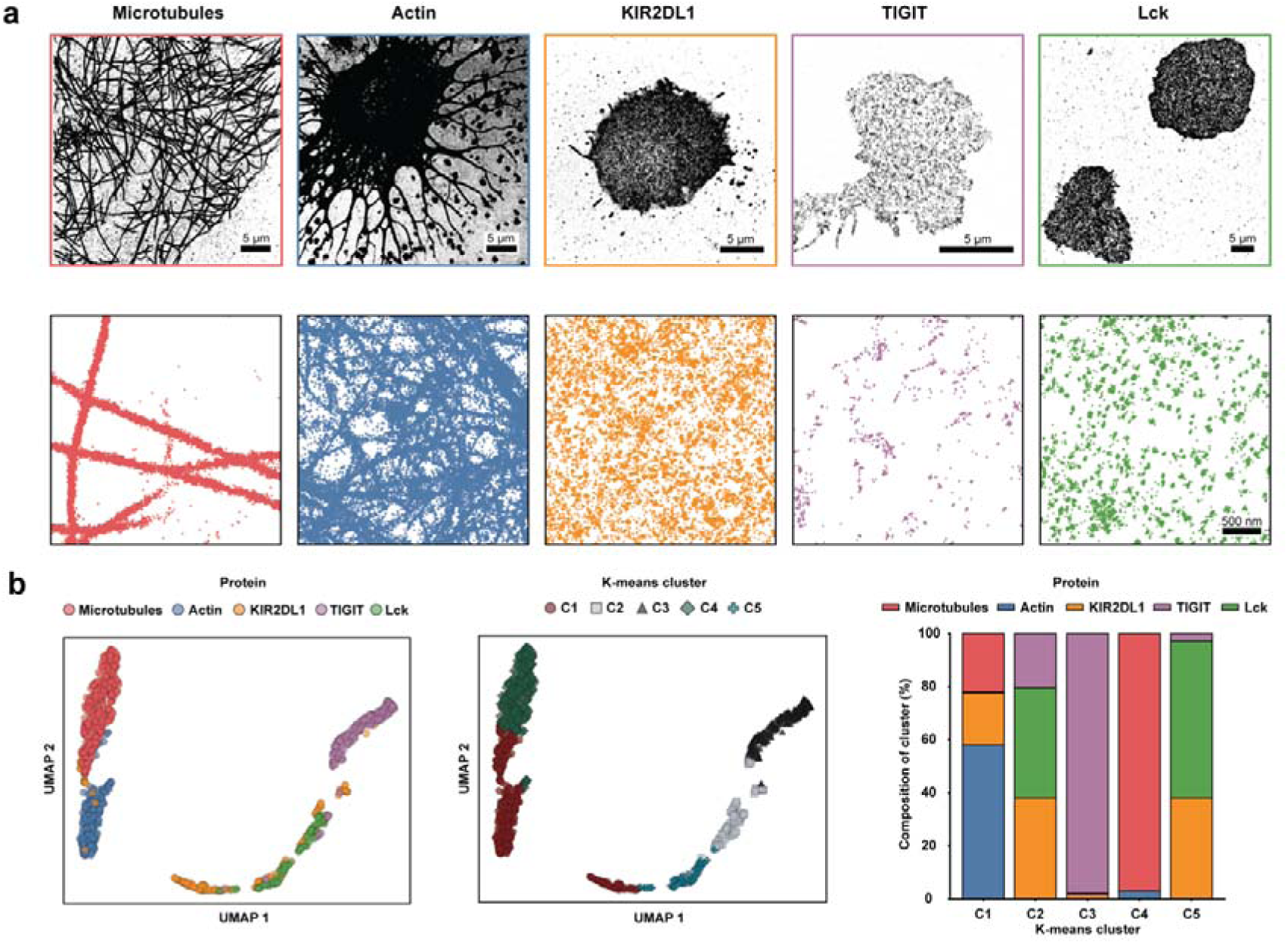
Contrastive learning distinguishes experimental protein distributions according to nanoscale organisation. **a)** Experimental SMLM datasets for microtubules (in COS7 cells), actin (in COS7 cells), KIR2DL1 (in NK cells), TIGIT (in primary CD8+ T cells) and Lck (in activated Jurkat E6.1 cells). For each protein, a representative full field of view (top) and example 3×3 µm ROI used for contrastive analysis (bottom) are shown. Scale bars, 5 µm for full fields of view and 500 nm for ROIs. **b)** ROIs were encoded into the 128D learned representation and visualised using UMAP, pseudo-coloured by protein identity (left) or K-means cluster assignment (centre). K-means was performed in the full 128D representation with k = 5. The percentage condition composition of each K-means cluster is shown (right). The analysis included 1,352 experimental ROIs: microtubules (n = 446), actin (n = 312), KIR2DL1 (n = 202), TIGIT (n = 267) and Lck (n = 125).

Because setting k equal to the number of experimental conditions imposes a known cluster number, we additionally estimated the preferred number of clusters using the elbow criterion without reference to protein identity (**Supplementary Figure S4**). This analysis selected k = 2, demonstrating that the dominant unsupervised structure of the representation is coarser than the five experimental labels and that the biological conditions do not necessarily correspond to five intrinsically discrete structural classes.

We next compared the learned representation directly with conventional spatial descriptors using the non-fibrous KIR2DL1, TIGIT and Lck datasets (**Supplementary Figure S5**). For each ROI, we calculated six predefined descriptors comprising DBSCAN cluster number, noise fraction, mean localisations per cluster and mean cluster area, together with Ripley-derived peak radius and maximum deviation. These classical descriptors contained substantial condition information: a repeated five-fold linear classifier achieved a balanced accuracy of 0.918 (95% range 0.869–0.955), compared with 0.956 (0.915–0.992) using the 128D contrastive representation. However, unsupervised K-means clustering showed clearer correspondence with protein identity in the learned representation, with ARI increasing from 0.398 (95% CI 0.338–0.464) for the six-dimensional classical descriptor space to 0.620 (95% CI 0.559–0.684) for the contrastive representation. Thus, predefined spatial descriptors can discriminate these proteins effectively, while the learned representation provides a more coherent unsupervised organisation without requiring selection of a particular set of spatial measurements.

Localisation density is itself a potentially informative feature of SMLM data but can also vary because of expression level, labelling efficiency or acquisition conditions. We therefore repeated the five-protein analysis after randomly subsampling ROIs to a fixed density of 100 localisations µm² (**Supplementary Figure S6**). Density normalisation retained structured separation between protein conditions and produced an ARI of 0.609 (95% CI 0.576–0.642; permutation p = 0.0002), indicating that the observed differences are not attributable solely to localisation number. Consistent with this, a separate systematic perturbation analysis of Lck ROIs showed little change after mild thinning, with progressively larger representation shifts only as localisation loss became severe (**Supplementary Figure S8**).

Next, we asked whether the learned representation was sensitive to biological context and to variation in experimental labelling (**Figure 4**). We first compared microtubule organisation across COS-7, HEK293 and HeLa cells, including Lck in Jurkat E6.1 cells as a structurally distinct contrast condition. Lck occupied a clearly separated region of the representation, whereas microtubule ROIs from the three adherent cell types showed substantial overlap but also condition-dependent enrichment within the shared manifold (**Figure 4a**). K-means clustering in the full 128D representation with k = 4 produced an ARI of 0.285 (95% CI 0.241–0.328; permutation p = 0.0002). Thus, cell type is associated with detectable differences in microtubule organisation, although these differences are considerably more subtle than those separating structurally distinct proteins. To illustrate the structural heterogeneity underlying this comparison, we mapped 52 microtubule ROIs from a representative COS-7 cell to their original cellular positions and corresponding locations in the learned representation (**Supplementary Figure S7**). Selected ROIs illustrate how variations in local microtubule organisation can be examined alongside their positions in the reference UMAP.

**Figure 4:**
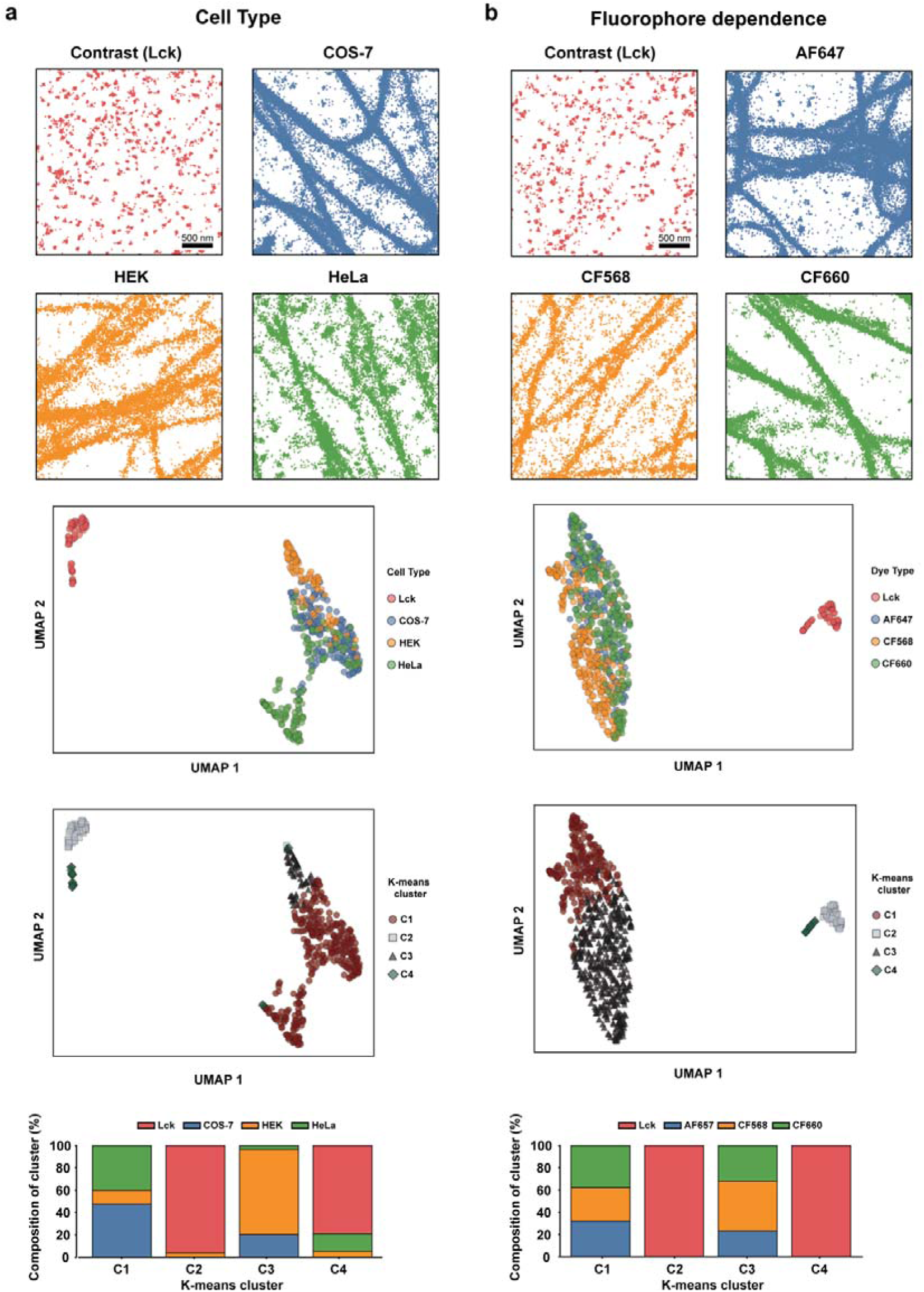
Contrastive learning reveals the relative effects of cell type and fluorophore choice on nanoscale organisation. **a)** Comparison of microtubule organisation across COS-7, HEK293 and HeLa cells, with Lck in Jurkat E6.1 cells included as a structurally distinct contrast condition. Representative 3×3 µm ROIs are shown above the condition-coloured UMAP, K-means-coloured UMAP and percentage cluster-composition plot. **b)** Comparison of microtubules labelled with AF647, CF568 or CF660, again including Lck as a structurally distinct contrast condition. For both analyses, K-means clustering was performed in the full 128D representation with k = 4; scale bars, 500 nm. The cell-type comparison included COS-7 (n = 184), HEK293 (n = 92), HeLa (n = 150) and Lck (n = 65) ROIs. The fluorophore comparison included AF647 (n = 184), CF568 (n = 276), CF660 (n = 238) and Lck (n = 65) ROIs.

We next compared microtubules labelled with AF647, CF568 or CF660, again including Lck as a structurally distinct contrast condition (**Figure 4b**). The three fluorophore conditions were extensively intermixed in the learned representation, while Lck remained clearly separated. Correspondingly, K-means clustering produced a lower ARI of 0.121 (95% CI 0.098–0.150; permutation p = 0.0002). These results indicate that, within this dataset, fluorophore choice contributes substantially less to the learned representation than protein identity and produces less structured variation than cell type.

Given the learned representations may also respond to technical variation in SMLM data, we systematically tested the effect of common acquisition and processing artefacts using 125 experimental Lck ROIs (**Supplementary Figure S8**). Increasing background, drift, localisation uncertainty, localisation loss and blinking/overcounting produced monotonic increases in paired representation-space displacement relative to the same unmodified ROI. The representation was comparatively insensitive to modest background contamination, small residual drift and mild thinning, whereas severe localisation loss, blur and drift produced substantially larger shifts. Combined multi-artifact perturbations likewise generated progressively greater displacement with increasing corruption severity. These results demonstrate that the learned representation can report graded technical degradation, while also emphasising the importance of consistent acquisition, processing and quality control when preparing and comparing SMLM datasets.

We next asked whether the learned representation could resolve graded pharmacological perturbation of a single nanoscale structure. COS-7 cells were treated with increasing concentrations of nocodazole, which progressively disrupts microtubule polymerisation and network organisation^25^ (**Figure 5a-b**). Representative ROIs showed increasing fragmentation of the microtubule network from untreated cells through 0.1 to 1.0 µg mL⁻¹ nocodazole. Consistent with this structural progression, the three treatment conditions occupied partially separated regions of the learned representation, with the highest-dose condition showing the clearest displacement from untreated cells. K-means clustering in the full 128D representation with k = 3 produced an ARI of 0.527 (95% CI 0.470–0.587; permutation p = 0.0002), demonstrating that the representation captures a graded drug-induced change in microtubule nanoscale organisation.

**Figure 5:**
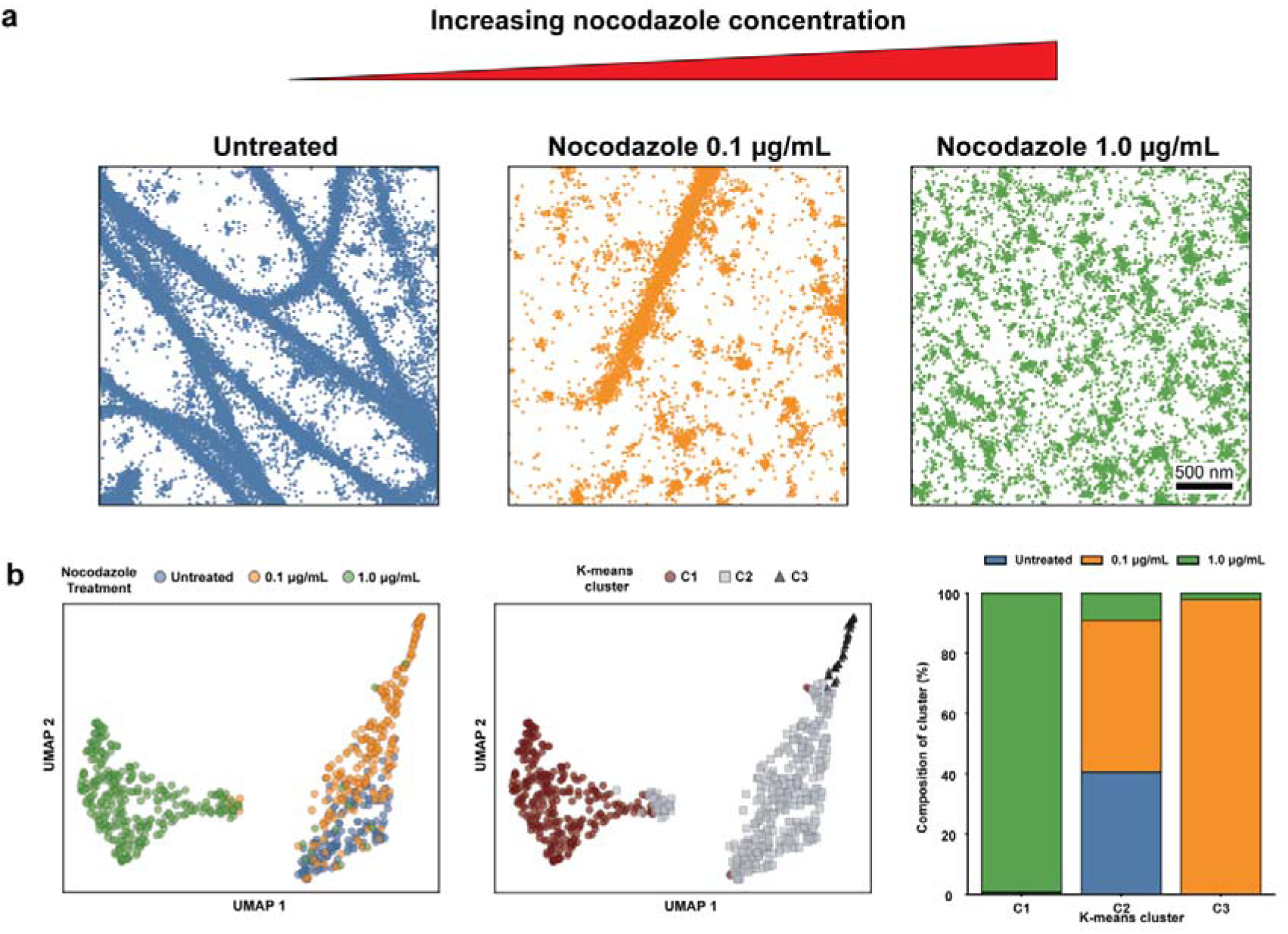
Contrastive learning resolves dose-dependent changes in microtubule nanoscale organisation following nocodazole treatment. **a)** Schematic illustrating progressive disruption of the microtubule network with increasing nocodazole concentration, together with representative 3×3 µm experimental ROIs from untreated COS-7 cells and cells treated with 0.1 or 1.0 µg mL⁻¹ nocodazole. Scale bar, 500 nm. **b)** Condition-coloured UMAP projection (left), K-means-coloured UMAP projection (centre) and percentage treatment composition of each K-means cluster (right). K-means clustering was performed in the full 128D representation with k = 3 and produced an ARI of 0.527 (95% CI 0.470–0.587). The analysis included untreated controls (n = 150), 0.1 µg/mL nocodazole (n = 229) and 1.0 µg/mL nocodazole (n = 330), giving 709 ROIs in total.

To determine whether the nocodazole response was driven primarily by treatment-dependent changes in localisation density, we repeated the analysis after subsampling all ROIs to 100 localisations µm⁻² (**Supplementary Figure S9a**). The treatment-dependent organisation remained evident after density normalisation, with an ARI of 0.561 (95% CI 0.505–0.620; permutation p = 0.0002). The similar clustering performance before and after density equalisation indicates that the nocodazole response is not attributable solely to localisation number.

We also applied the framework to two additional biological perturbation datasets (**Supplementary Figure S9b-c**). Actin ROIs from early and late Jurkat E6.1 immunological synapses, sampled from central and peripheral regions, showed partial separation between conditions, with K-means clustering giving an ARI of 0.227 (95% CI 0.130–0.385; permutation p = 0.0002). By contrast, EGFR ROIs from resting HeLa cells and cells stimulated with EGF for 5 or 15 min showed extensive overlap, with a lower ARI of 0.115 (95% CI 0.090–0.147; permutation p = 0.0002). These datasets demonstrate that correspondence between the learned representation and experimental condition varies across biological systems, with some perturbations producing substantial overlap. The framework therefore provides a common basis for comparing organisational variation but does not imply that every experimental condition corresponds to a distinct nanoscale structural state.

## Discussion

We present a self-supervised contrastive learning framework that enables SMLM point patterns to be analysed within a shared representation of nanoscale spatial organisation. By learning from procedurally generated spatial patterns without biological class labels, the approach provides a common feature space in which heterogeneous experimental datasets can be compared without specifying the spatial characteristics of interest in advance (**Figure 1**). Across simulated and experimental data, the learned representation captured differences in clustering, density and fibre-like organisation, distinguished proteins with different nanoscale architectures and resolved graded changes following pharmacological perturbation (**Figures 2-5**). In particular, the progressive disruption of microtubules by nocodazole was reflected in the learned representation, even after normalisation of localisation density (**Figure 5** and **Supplementary Figure S9**). These results establish contrastive representation learning as a flexible approach for exploratory nanoscale phenotyping, thereby extending SMLM analysis beyond comparisons based on individual, predefined spatial measurements.

A principal advantage of this approach is the ability to investigate complex spatial variation using a common multivariate representation. Established methods, such as DBSCAN and Ripley’s K-function, provide valuable and physically interpretable measurements^14^, but their application requires choices about which aspects of spatial organisation to quantify. Our direct benchmark demonstrated that six conventional spatial descriptors retained substantial discriminatory information, achieving high supervised classification performance for KIR2DL1, TIGIT and Lck. Nevertheless, the contrastive representation produced substantially better agreement with protein identity during unsupervised clustering (ARI = 0.620 versus 0.398). This distinction is important for exploratory experiments, where the relevant structural differences may not be known beforehand. Rather than selecting clustering or spatial-statistical measurements at the outset, researchers can use a common learned representation to identify relationships between experimental conditions, detect structural variation and prioritise subsequent investigations. Conventional spatial statistics can then be applied to characterise the specific features underlying these differences.

The broader opportunity lies in using learned spatial representations to characterise biological heterogeneity at scales that are difficult to investigate systematically using conventional SMLM analysis. For example, large collections of ROIs from individual cells could be combined to develop cell-level nanoscale phenotypes, enabling comparisons across cell populations, differentiation states or pharmacological responses. Applied to primary cells or heterogeneous tissues, such analyses could help identify previously unrecognised organisational states or rare cellular subpopulations that are not readily distinguished by conventional markers. Similarly, screening experimental ROIs for unusual representations could reveal rare structural phenotypes, heterogeneous responses or anomalous samples for further investigation. These applications would require appropriate cell-level aggregation, experimental replication and independent validation, but illustrate how representation learning could shift SMLM towards discovery-led analysis rather than exclusively hypothesis-driven measurements. Combining representations from multiple molecular targets, or integrating them with signalling, functional and other single-cell measurements, offers a further opportunity to investigate how nanoscale organisation relates to cellular identity and function.

An important consideration for such applications is the distinction between biological variation and technical differences in SMLM data^8^. Our density-normalisation experiments showed that differences between protein architectures and the nocodazole response were not explained solely by localisation number. Systematic perturbations of experimental Lck datasets further demonstrated that the representation responds predictably to increasing localisation uncertainty, drift, thinning, background contamination and overcounting. These findings suggest that the learned representation could provide an additional readout of dataset consistency, but not a replacement for established SMLM quality-control measures, *i.e.*, localisation precision assessment, drift correction and evaluation of background and repeated blinking. At the same time, the response of the representation depends on the range of spatial patterns encountered during training. Analysis of the Perlin training distribution demonstrated substantial, but incomplete, coverage of experimental protein architectures, with lower representation-space support for actin and microtubule datasets. Controlled simulations also revealed saturation at extreme fibre densities, illustrating that distances in the learned representation cannot be assumed to scale linearly with underlying physical changes. Broader synthetic training distributions, additional experimental reference data and systematic evaluation across acquisition platforms will therefore be important steps towards improving transferability and establishing the reliability of future applications. Although the present implementation is restricted to two-dimensional SMLM data, extension to 3D is technically feasible through volumetric localisation histograms and three-dimensional convolutional encoders. Such an extension would require additional computational resources, appropriate 3D training distributions and consideration of anisotropic localisation precision, particularly the typically lower precision of axial measurements^26^.

Ultimately, we envisage nanoscale spatial-omics developing as a complementary layer of cellular phenotyping alongside established molecular and functional measurements. A central requirement will be the availability of large, diverse and well-annotated SMLM datasets that allow spatial representations to be evaluated and compared across experiments and laboratories. Community resources such as nano-org^22^ provide a foundation for this development, connecting reusable experimental datasets with computational methods for comparative analysis. Coupled with standardised acquisition, metadata and quality-control practices, representation-learning frameworks could support the construction of reference maps of nanoscale organisation, systematic comparisons of biological diversity and the identification of previously unrecognised structural phenotypes. The approach presented here establishes a foundation for this direction, providing a common representation through which nanoscale organisation can be explored, compared and ultimately connected to biological function.

## Methods

### Contrastive learning

Contrastive representation learning was implemented in PyTorch using an adapted SimCLR-style framework^19^. Single-molecule localisation coordinates within each 3×3 µm region of interest (ROI) were rasterised as a single-channel 120×120 localisation-count histogram, corresponding to a spatial sampling of 25 nm per bin. Histograms were normalised by their total localisation count before input to the network. The 25 nm bin size defines the spatial sampling of the encoder input rather than the experimentally achieved localisation precision or spatial resolution. For applications examining substantially different structural length scales or imaging modalities, the physical ROI dimensions and histogram bin size can be adjusted, although changes to these parameters would require appropriate model training or validation to establish performance at the new spatial scale.

The final encoder architecture (“Pool8”) comprised five convolutional layers with channel dimensions 1>8>16>32>32>32. All convolutions used 3×3 kernels with padding and were followed by batch normalisation and Mish activation^27^. Two 2×2 average-pooling stages were used for spatial downsampling, and Dropout2D (p = 0.2) was applied within the first 32-channel convolutional block. Following the final convolutional layer, adaptive average pooling generated a 32×8×8 representation, corresponding to 2048 features. These were flattened and projected through a fully connected layer to a 128D encoder representation, followed by dropout (p = 0.2), layer normalisation and Mish activation. During training, the encoder was followed by a projection head comprising three fully connected layers (128>128>128>512), with dropout, layer normalisation and Mish activation in the intermediate layers and a sigmoid activation at the final output. The complete network contained 386,736 trainable parameters, of which 287,152 belonged to the encoder. The projection head was discarded after training, and all downstream analyses used the L2-normalised 128D encoder representation.

Training point patterns were generated procedurally using fractal Perlin noise^24^ with the FractalPerlin2D implementation from the pyperlin package. Perlin noise was used as a flexible procedural training distribution rather than as a mechanistic model of any particular protein architecture: varying its generative parameters produces correlated spatial patterns spanning multiple spatial scales without requiring biological class labels or specification of discrete morphologies. Each simulated pattern represented a 3×3 µm field and contained between 300 and 9999 signal localisations (approximately 33–1111 localisations µm⁻²).

Generative parameters varied the spatial scale of the Perlin field, the range and weighting of multiscale components, localisation number and whether localisations were selected from the upper or lower region of the generated noise field. Specifically, the field-size exponent was sampled from 7–12, the multiscale parameter from 5 to one below the field-size exponent, and the scale-weighting parameter from 0–3.

Two independently generated patterns sharing the same generative parameter vector were treated as a positive pair. Technical variation was introduced independently into each pattern during generation. Uniform background localisations corresponding to 0–10% of the signal localisation number were added, 0–15% of signal localisations were randomly removed, and localisation coordinates were displaced by Gaussian noise whose per-localisation standard deviation was sampled from a gamma distribution with mean 20 nm. Thus, the synthetic training distribution did not explicitly model a foreground biological structure plus a fixed background; rather, correlated spatial organisation was generated procedurally and uniform background contamination was introduced separately.

The final training bank comprised 8000 parameter groups generated with seed 1, with two independently generated positive views stored for each group. A separate validation bank comprised 1000 parameter groups generated with seed 2. Each training epoch traversed all 8000 training groups. For each optimisation step, the two positive images were contrasted against 15 negative parameter groups, each represented by two independently generated images, giving 32 images per contrastive step. We used an adapted NT-Xent-style objective^19^ with cosine similarity and temperature τ = 0.1. Negative similarities were weighted according to the summed difference between normalised generative parameter vectors, such that negative examples generated from more different parameter settings contributed more strongly to the loss, while closely related parameter settings were penalised less strongly.

Optimisation used AdamW^28^ with an initial learning rate of 3×10^−^□ and weight decay of 1×10^−^□. Learning rate was reduced on plateau by a factor of 0.5 (scheduler patience = 1; minimum learning rate = 1×10^−^□). Training proceeded for a maximum of 100 epochs with early-stopping patience of three epochs. The final model was trained on a single NVIDIA A100 GPU and required approximately 15 min. Three independently trained Pool8 models were evaluated across five simulated benchmark families, yielding mean adjusted Rand indices of 0.845, 0.827 and 0.785 for seeds 1, 2 and 3, respectively. The seed-1 checkpoint was selected based on its aggregate simulated benchmark performance and used for the experimental analyses reported in the main manuscript.

### Training-distribution coverage

To assess the extent to which experimental SMLM data were represented within the synthetic training distribution, one 128D encoder representation was generated for each of the 8000 Perlin training parameter groups and for each of the 1,000 independently generated validation groups. Training support was quantified directly in the L2-normalised 128D representation using cosine distance. For each training embedding, the cosine distance to its tenth-nearest other training embedding was calculated, and the 95th percentile of this distribution was used as a training-support threshold. For each experimental ROI or independently generated Perlin validation sample, a training-support ratio was calculated as:

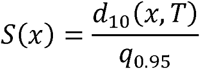

where *d*_10_(*x,y*), is the cosine distance from sample *x* to its tenth-nearest training representation in *T*, and *q*_0.95_. is the 95th percentile of the corresponding tenth-nearest-neighbour distances within the training set, excluding self-matches. The tenth-nearest neighbour was selected to provide a measure of local representation-space coverage that is less sensitive to isolated close matches than the single nearest-neighbour distance. Samples with *S*(*x*)≤1 were classified as falling within training support.

Each experimental ROI from **Figure 3** and each independently generated Perlin validation sample was assigned a training-support ratio, *S*(*x*). A joint UMAP of training and experimental embeddings was generated for visualisation only (**Supplementary Figure S1b**).

### Analysis

SMLM ROIs were represented by L2-normalised 128D encoder vectors. Unless otherwise stated, K-means clustering was performed directly in this full 128D representation rather than on a dimensionality-reduced projection. The number of clusters, *k*, was set to the number of experimental or simulated conditions for condition-composition analyses. K-means was initialised 10 times and a fixed random seed was used for reproducibility. Agreement between unsupervised cluster assignments and known condition labels was quantified using the adjusted Rand index (ARI)^29^.

ARI confidence intervals were estimated by nonparametric bootstrapping. For each of 1,000 bootstrap replicates, paired condition and cluster labels were sampled with replacement and the ARI recalculated; the 2.5^th^ and 97.5^th^ percentiles of the resulting distribution were used as the 95% confidence interval. Statistical significance was assessed by randomly permuting experimental-condition labels 5,000 times while holding K-means cluster assignments fixed. ARI was recalculated for each permutation, and a one-sided empirical P value was calculated as follows, where *n_extreme_* denotes the number of permutations producing an ARI greater than or equal to the observed ARI;

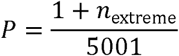

This procedure gives a minimum attainable *P* value of approximately 0.0002.

For **Supplementary Figure S4**, the number of K-means clusters was additionally estimated without reference to experimental labels using the elbow method. Within-cluster sum of squares was calculated for successive values of k in the full 128D representation, and the elbow was identified from the curvature of this relationship.

For visualisation only, 128D representations were projected into two dimensions using Uniform Manifold Approximation and Projection (UMAP)^21^ with *n_neighbors* = 15, *min_dist* = 0.1, cosine distance and a fixed random seed. K-means clustering, ARI calculation and other quantitative comparisons were not performed on the UMAP coordinates.

For the direct comparison with conventional SMLM descriptors, KIR2DL1, TIGIT and Lck ROIs were analysed independently of the contrastive representation. DBSCAN^30^ was applied to localisation coordinates using ε = 25 nm and minPts = 5. Four descriptors were extracted for each ROI: number of clusters, fraction of localisations classified as noise, mean number of localisations per cluster and mean cluster area, with cluster area defined from the two-dimensional convex hull of clustered localisations. Ripley analysis^31^ was additionally performed over radii from 5 to 300 nm in 5-nm increments. Ripley’s K-function was transformed into the H-function, 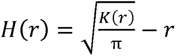, from which two descriptors were extracted: the radius at which H(r) reached its maximum and the corresponding maximum H(r) value. The six resulting descriptors were imputed where necessary and standardised to zero mean and unit variance to form a six-dimensional classical feature representation.

To compare the classical and learned representations directly, K-means clustering with k = 3 was independently applied to the standardised six-dimensional classical representation and the 128D contrastive representation, and correspondence with protein identity was quantified by ARI with 1,000-replicate bootstrap confidence intervals. Condition discriminability was additionally evaluated using multinomial linear logistic regression with class weighting. Balanced accuracy was estimated using five repeats of stratified five-fold cross-validation, with individual ROIs assigned to training and test folds and stratification performed by protein identity. Identical cross-validation splits and evaluation procedures were applied to the classical and contrastive representations.

To assess dependence on localisation density, experimental ROIs were randomly subsampled to a fixed density of 100 localisations µm⁻², corresponding to 900 localisations within each 3×3 µm ROI. Sampling was performed without replacement; ROIs containing fewer than the required number of localisations were excluded from the density-normalised analysis. The thinned ROIs were then passed through the same encoder and downstream analysis pipeline as the unthinned data. Density-normalised analyses were performed for the five-protein comparison and the nocodazole dose-response dataset.

To quantify sensitivity to common SMLM artefacts, 125 experimental Lck ROIs were subjected to controlled in silico perturbations and compared with the corresponding unmodified ROI. Perturbations comprised random thinning retaining 90, 75, 50, 25 or 10% of localisations; additional Gaussian localisation blur with standard deviations of 5, 10, 20, 30 or 50 nm per coordinate; uniform Poisson background at 1, 2.5, 5, 10, 20 or 50 localisations µm⁻²; linear drift with total first-to-last displacements of 10, 25, 50, 100 or 200 nm; and simulated blinking/overcounting producing expected localisation-number multipliers of 1.25, 1.5, 2, 3 or 5, with repeated localisations displaced by Gaussian jitter of 10 nm. Additional mild, moderate and severe multi-artifact conditions combined several perturbations. Each corrupted ROI was encoded independently and its displacement from the matched unmodified ROI quantified using cosine distance in the 128D representation. Confidence intervals for median paired cosine distances were obtained by bootstrap resampling of source ROIs. Monotonicity of representation displacement with artefact severity was assessed within ROIs using Spearman rank correlation.

### Data Simulations

Independent simulated datasets were generated to evaluate how the learned representation responded to controlled variation in nanoscale point-pattern organisation. These simulations were distinct from the Perlin-noise simulations used to train the encoder. All evaluation simulations were generated within 3×3 µm ROIs with the simulation parameters collated in **Table 1**.

For Gaussian-cluster simulations, cluster centres were positioned randomly within the ROI and localisations associated with each cluster were sampled from two-dimensional isotropic Gaussian distributions. Simulation families independently varied the number of clusters, number of localisations per cluster, Gaussian standard deviation and amount of uniformly distributed background. Cluster-number simulations used 2, 5, 9, 10 or 20 Gaussian clusters. Additional parameter sweeps used 10, 50, 90, 100 or 200 localisations per cluster; Gaussian standard deviations of 30, 35, 40, 49, 50, 100 or 200 nm; and background-density parameters of 0.1, 0.4, 0.5 or 0.9. Parameters not under investigation were held constant within each simulation family.

To model mixtures of clustered and fibre-like nanoscale organisation, ROIs contained ten structures comprising five isotropic Gaussian clusters and five elongated Gaussian structures. Gaussian clusters contained 100 localisations with a standard deviation of 50 nm. Fibre-like structures were generated from anisotropic two-dimensional Gaussian distributions with a major-axis standard deviation of 1000 nm and minor-axis standard deviation of 50 nm and were assigned a random orientation. A fixed background of 1000 uniformly distributed localisations was included. Fibre density was varied by changing the number of localisations in each elongated structure relative to the 100 localisations per Gaussian cluster. The principal simulation series used relative fibre densities of 1, 2, 3, 5, 10 and 20x.

To examine saturation of the learned representation at extreme structural densities, the fibre-density series was extended to 1, 2, 3, 5, 10, 20, 50, 100, 200 and 500x, with 30 independently simulated ROIs generated at each level. Points generated outside the 3×3 µm ROI were discarded. Representation change was quantified as the cosine distance between each ROI and the 1x reference distribution in the full 128D representation, with bootstrap confidence intervals calculated across simulated ROIs.

### Software

All computational analyses were performed in Python. Neural-network training and inference were implemented using PyTorch^32^. Numerical and tabular analyses used NumPy and pandas; K-means clustering, DBSCAN, logistic-regression classification and associated metrics used scikit-learn^33^; UMAP projections were generated using umap-learn^21^; and statistical and geometric calculations used SciPy where appropriate. Analysis scripts, trained model checkpoints and code required to reproduce the analyses are provided with the accompanying software repository (https://gitlab.bham.ac.uk/owendz-protein-databank/smlm-contrastive-learning).

### Experimental methods

dSTORM microtubule data in **Figures 4**, **5** and **Supplementary Figures S1, S4, S7** and **S9** as well as Lck data (**Figures 3** and **4**; **Supplementary Figure S4, S5, S6** and **S8**) were acquired in-house. DNA-PAINT EGFR data in **Supplementary Figure S9** was kindly provided by Alexandra Kaminer from the Heilemann lab. Experimental methods for these datasets are described below, all other experimental data is available on nano-org, with experimental methods available through their DOIs.

### dSTORM experimental data

#### Cell culture

COS-7, HeLa, and HEK293 cells were cultured in Dulbecco’s Modified Eagle Medium (DMEM, high glucose; Sigma-Aldrich), supplemented with 10% fetal bovine serum (FBS; Gibco, Life Technologies), 1% penicillin–streptomycin (Gibco, Life Technologies), and 1% L-glutamine (Gibco, Life Technologies). Jurkat E6.1 cells were cultured in RPMI 1640 (Sigma-Aldrich) with the same supplement concentrations. All cells were maintained at 37 °C in a humidified incubator with 5% CO. For imaging, adherent cells were seeded at a density of 1×10 cells per well into eight-well µ-slides (Ibidi, glass bottom) one day prior to fixation. HEK wells were pre-treated with fibronectin (Sigma-Aldrich) (1:100 dilution in phosphate-buffered saline (PBS); Gibco) for 30 minutes before cell seeding. Jurkat cells were seeded at 2×10 cells per chamber in six-channel µ-slides (Ibidi, glass bottom) pre-coated with anti-CD3/CD28 (1μg/ml; Invitrogen). Cells were activated at 37 °C for 5 minutes, then fixed in 4% paraformaldehyde (PFA; Sigma-Aldrich) for 10 minutes and washed three times with PBS. These samples were used for Lck imaging.

#### Sample preparation

For microtubule disruption experiments, COS-7, HeLa, and HEK cells were incubated in DMEM containing 0.1 µg/mL or 1 µg/mL nocodazole (Tocris) for 30 minutes at 37 °C. Cells were then washed three times with PBS. Untreated controls were PBS-washed directly. To preserve microtubule architecture, cells were subjected to sequential extraction and fixation. Cells were first extracted for 90 seconds at 37 °C using a pre-warmed solution of 0.25% Triton X-100 (Sigma-Aldrich) and 0.1% glutaraldehyde (Sigma-Aldrich) in PEM buffer (80 mM PIPES (Bioworld), 5 mM EGTA (ThermoFisher Scientific), 2 mM MgCl (Invitrogen), pH 6.8; (Invitrogen)), followed by fixation for 10 minutes at 37 °C in 0.25% Triton X-100 and 0.5% glutaraldehyde in PEM. After fixation, cells were quenched with 1 mg/mL sodium borohydride (Sigma-Aldrich) for 7 minutes and washed three times with PBS. All SMLM imaging was performed using an ONI Nanoimager S microscope unless otherwise stated.

#### dSTORM imaging

Cells were permeabilised with 0.1% Triton X-100 in PBS for 3 minutes at room temperature and blocked with 5% bovine serum albumin (BSA; Sigma-Aldrich) for 30 minutes. For microtubule staining, cells were incubated with mouse monoclonal anti-β-tubulin IgG (200 µg/mL; Santa Cruz Biotechnology), diluted 1:50 in 5% BSA, for 30 minutes at room temperature. After three PBS washes, cells were incubated in Alexa Fluor™ 647-conjugated goat anti-mouse IgG (2 mg/mL; Life Technologies), diluted to 2 μg/ml in 5% BSA, for 30 minutes in the dark. If specified, CF568 and CF660 conjugated goat anti-mouse IgG (2 mg/mL; Sigma-Aldrich) were used under the same conditions.

For Lck staining, cells were incubated with 1μg/ml Lck (D88) XP® Rabbit mAb (Cell Signalling Technology) in 5% BSA, for 30 minutes at room temperature. After washing three times with PBS, samples were incubated in Alexa Fluor™ 647-conjugated goat anti-rabbit IgG (2 mg/mL; Life Technologies), diluted 1:1000 in 5% (w/v) BSA, for 30 minutes in the dark. Finally, all samples were washed ten times with PBS. Immediately before dSTORM imaging, PBS was replaced with an imaging buffer containing 18% glucose, 10 mM Tris (pH 8), 50 mM NaCl, 0.8 mg/mL glucose oxidase, 50 mM cysteamine, and 40 µg/mL catalase (all from Sigma-Aldrich).

#### Data analysis

Data analysis was conducted using the Super resolution Microscopy Analysis Platform (SMAP)^34^, with default settings applied unless stated otherwise. Single-molecule localisations were fitted using the *PSF free* algorithm. Localisations with an estimated precision >30 nm were excluded, consistent with common SMLM practice to remove low-confidence points and ensure reliable nanoscale resolution. Drift correction and grouping were performed to mitigate the effects of sample drift and multiple blinking of fluorophores. ROIs were generated from full field-of-view localisation files by manually defining polygons that closely followed the outer contour of each cell. The regions enclosed by these polygons were subsequently divided into a regular grid of 3×3 µm ROIs for analysis, thereby restricting sampling to the cellular region.

### DNA-PAINT experimental data

#### Cell culture

The human cervical cancer cell line HeLa (# ACC 57, DSMZ, Braunschweig, Germany) was cultured in Dulbecco’s modified Eagle’s medium (DMEM) (# 11574486, Gibco, Life Technologies, Waltham, MA, USA) supplemented with 10% fetal bovine serum (# 35-079-CV, Corning Inc., Corning, NY, USA), 1 unit/mL penicillin and 1 µg/mL streptomycin (Gibco, Life Technologies) and 1% v/v GlutaMAX (# 35050-038, Gibco, Life Technologies). The cells were incubated at 37 °C with 5% CO_2_ and were passaged every 3–4 days. HeLa cells were seeded in ibidi µ-Slide (# 80607, ibidi GmbH, Gräfelfing, Germany) coated with PLL-PEG-RGD (poly-l-lysine-grafted polyethylene glycol modified with a CGRGDS peptide) at the density of 1×10^5 cells/mL for one day growth.

#### Sample preparation

The cells were starved for 2 hours prior stimulation using the introduced growth media without fetal bovine serum. After starvation, EGF (# AF-100-15, PeproTech, Thermo Fisher Scientific, Waltham, MA, USA) was diluted in the serum free media to end concentration of 100 ng/mL and then incubated for either 5 or 15 min. Both, resting and stimulated cells were fixed using 3% formaldehyde (FA) (# 28908, Sigma-Aldrich, St. Louis, MO, USA) with 0.25% glutaraldehyde (GA) (# G5882, Sigma) and incubated for 15 min at 37 °C. For EGFR labeling, a monoclonal primary antibody (1:50 dilution, #sc-120, Santa Cruz Biotechnology, Texas, USA) was pre-incubated with a secondary nanobody (2-fold excess) conjugated to the R3 docking strand (Massive Photonics, München, Germany) for 1 h at 4 °C in the blocking buffer (1 mM EDTA, 0.02% Tween 20, 0.05% NaN3, 2% bovine serum albumin (BSA, # 9048-46-8, Carl Roth GmbH & Co. KG, Karlsruhe, Germany), 0.05 mg/mL salmon sperm DNA in PBS). Fixed cells were blocked in the blocking buffer for 20 min and incubated with the pre-incubated antibody/nanobody mixture for 2 h at room temperature. Gold beads (# A11-100-NPC-DIH-1-100, Nanopartz Inc., Loveland, US) were diluted to 1:5 in PBS and added to each well for 15 min, followed by washing with PBS. The cells were post-fixed using 4% FA.

#### DNA-PAINT imaging

DNA-PAINT imaging was performed on a home-built widefield setup based on a custom-built widefield microscope based on a Nikon Eclipse Ti inverted microscope. Excitation was provided by a 561 nm laser (200 mW Sapphire, Coherent Inc., Santa Clara, CA, USA) with laser power modulated via an acousto-optic tunable filter (AOTFnC-400.650-TN, AA Opto Electronic, France). To ensure a clean beam profile, the laser was fiber-coupled using a collimator (60FC-4-M6.2-33) into a polarization-maintaining single-mode optical fiber (PMC-E-400RGB), and then re-collimated to a 6 mm full width at half maximum (FWHM) beam (60FC-T-4-M50L-01; all from Schäfter & Kirchhoff GmbH, Germany). The collimated beam was expanded using a telescope (AC255-030-A-ML and AC508-150-A-ML, Thorlabs GmbH, Germany) and focused onto the back focal plane of a 100x TIRF oil immersion objective (CFI Apochromat TIRF 100XC Oil, Nikon, Japan). A motorized mirror (MTS50-Z8, Thorlabs) enabled adjustment of the illumination angle for widefield, HILO, or TIRF imaging modes. Axial focus was stabilized using Perfect Focus System (Ti-PFS, Nikon), while lateral sample positioning was controlled by a motorized stage (Ti-S-ER, Nikon) in combination with a piezo stage (Nano-Drive, MadCityLabs, USA). Excitation light was introduced into the microscope via a multiband dielectric beamsplitter (zt405/488/561/640rpc, AHF Analysentechnik, Germany), which also directed emission light into the detection path. Fluorescence was spectrally filtered with a bandpass filter (610/60 ET, Chroma) and imaged with an EMCCD camera (iXon Ultra DU-897U-CS0, Andor, Northern Ireland).

Prior to imaging, the R3 imager strand (conjugated to Cy3B; Massive Photonics) was diluted to a final concentration of 2 nM in an imaging buffer containing 5 mM Tris/HCl (pH 8.0), 75 mM MgCl_2_, and 0.05% Tween-20, freshly supplemented with an oxygen scavenging and triplet-state quenching system (1xPCA, 1xPCD, and 1xTrolox), following a published protocol ^35^. DNA-PAINT imaging was conducted using a 561 nm laser at an excitation intensity of 0.1 kW/cm^2^. All microscope components were controlled via the µManager software^36^. Image stacks of 20,000 frames were recorded with the following parameters: 100 ms exposure time, EM gain of 150, 3x preamplifier gain, 10 MHz readout rate, image size of 256×256 pixels, and frame transfer mode activated. Cell positions were saved within µManager to facilitate subsequent multi-target imaging. Bright-field images were acquired before and after each DNA-PAINT acquisition.

#### Data analysis

Image processing was performed using Picasso software (https://github.com/jungmannlab/Picasso). First, single emitters in each frame were localized by fitting the Maximum Likelihood Estimation for Integrated Gaussian parameters. Next, drift correction was performed using fiducial markers. Localized single molecule events were filtered for width of the point spread function (PSF), localizations which appeared in multiple consecutive frames were merged with parameters based on the NeNA (nearest neighbor based analysis) value which represents experimental localization precision: radius of 1 times NeNA and 4 min. dark frame. Next DBSCAN (density-based spatial clustering and application with noise) clustering was performed using 1xNeNA and 12min.sample. The identified clusters were further filtered based on the mean frame time within a range of µ-2*δ* to µ+2*δ*, where µ represents the average mean frame time and *δ* is the standard deviation, and the *δ* within a range from 1500 to 8000 frames.

## Acknowledgements

Data was acquired in the COMPARE Advanced Imaging Facility at the University of Birmingham. DMO and SS acknowledge funding from the Biotechnology and Biological Sciences research Council (BBSRC) grant BB/X018644/1. RH received funding from the European Research Council (ERC) through grant 101001332-SelfDriving4DSR and Horizon Europe through grants 101057970-AI4LIFE and 101099654-RTSuperES. Views and opinions expressed are, however, those of the authors only and do not necessarily reflect those of the European Union. Neither the European Union nor the granting authority can be held responsible for them. This work was also supported by a European Molecular Biology Organization (EMBO) installation grant (EMBO-2020-IG-4734 to RH). MH acknowledges funding from Collaborative Research Centre (CRC) 1507 and the Interfacing image analysis and molecular life science (iMOL) initiative.

## Author Contributions

DJN, LGJ, SS, and MCW developed the contrastive learning framework. DJN, SS, MCW, AK, KS, MH and RP provided simulations, experimental data and testing. LGJ, PR-D, RH, SFL, DJN and DMO conceived the work. DJN and DMO wrote the manuscript.

## Code and data availability

All experimental data is stored and available for download on https://nano-org.bham.ac.uk. The source code for the contrastive learning framework and associated analysis workflows is available through the project GitLab repository (https://gitlab.bham.ac.uk/owendz-protein-databank/smlm-contrastive-learning). An archived version of the software is also available on Zenodo (https://doi.org/10.5281/zenodo.24079915).

## Notes

https://nano-org.bham.ac.uk

https://gitlab.bham.ac.uk/owendz-protein-databank/smlm-contrastive-learning

### Competing Interest Statement

The authors have declared no competing interest.

### Summary of Updates

Model architecture: Optimised and retrained the contrastive learning model using the revised Pool8 encoder architecture, with improved feature extraction and a 128-dimensional learned representation. Training and documentation: Expanded the description of synthetic Perlin-noise training data, network architecture, training parameters and preprocessing procedures. Simulation analyses: Extended simulations to investigate structural variation, relationships between simulation families and saturation of representation responses at extreme densities. Training-distribution coverage: Added quantitative analyses assessing how well the synthetic training distribution represents experimental SMLM point patterns. Conventional methods benchmark: Introduced a direct comparison against six DBSCAN- and Ripley-derived spatial descriptors, evaluating unsupervised clustering and supervised classification. Experimental reanalysis: Reprocessed experimental datasets using the revised encoder and updated the quantitative clustering results. Density and artefact controls: Expanded localisation-density normalisation experiments and introduced systematic assessments of drift, localisation uncertainty, background contamination, thinning and overcounting. Biological interpretation: Added a cell-linked microtubule visualisation and refined the interpretation of experimental differences and methodological limitations. Manuscript and figures: Substantially revised the Introduction, Results, Discussion and Methods; updated all five main figures and expanded the Supplementary Information from four to nine figures.

## References

1 Visscher, P. M. et al. 10 Years of GWAS Discovery: Biology, Function, and Translation. Am J Hum Genet 101, 5–22, doi:10.1016/j.ajhg.2017.06.005 (2017).

2 Genomes Project, C., et al. A global reference for human genetic variation. Nature 526, 68–74, doi:10.1038/nature15393 (2015).

3 Tang, F. et al. mRNA-Seq whole-transcriptome analysis of a single cell. Nature Methods 6, 377–382, doi:10.1038/nmeth.1315 (2009).

4 Tabula Muris, C. Single-cell transcriptomics of 20 mouse organs creates a Tabula Muris. Nature 562, 367–372, doi:10.1038/s41586-018-0590-4 (2018).

5 Jumper, J. et al. Highly accurate protein structure prediction with AlphaFold. Nature 596, 583–589, doi:10.1038/s41586-021-03819-2 (2021).

6 Ståhl, P. L. et al. Visualization and analysis of gene expression in tissue sections by spatial transcriptomics. Science 353, 78–82, doi:10.1126/science.aaf2403 (2016).

7 Goltsev, Y. et al. Deep Profiling of Mouse Splenic Architecture with CODEX Multiplexed Imaging. Cell 174, 968–981.e915, doi:10.1016/j.cell.2018.07.010 (2018).

8 Lelek, M. et al. Single-molecule localization microscopy. Nat Rev Methods Primers 1, 39, doi:10.1038/s43586-021-00038-x (2021).

9 Balzarotti, F. et al. Nanometer resolution imaging and tracking of fluorescent molecules with minimal photon fluxes. Science 355, 606–612, doi:10.1126/science.aak9913 (2017).

10 Zhou, Y. et al. Membrane potential modulates plasma membrane phospholipid dynamics and K-Ras signaling. Science 349, 873–876, doi:10.1126/science.aaa5619 (2015).

11 Truong Quang, B. A., et al. Extent of myosin penetration within the actin cortex regulates cell surface mechanics. Nature Communications 12, 6511, doi:10.1038/s41467-021-26611-2 (2021).

12 Sengupta, P. et al. Probing protein heterogeneity in the plasma membrane using PALM and pair correlation analysis. Nat Methods 8, 969–975, doi:10.1038/nmeth.1704 (2011).

13 Nieves, D. J. & Owen, D. M. Analysis methods for interrogating spatial organisation of single molecule localisation microscopy data. Int J Biochem Cell Biol 123, 105749, doi:10.1016/j.biocel.2020.105749 (2020).

14 Nieves, D. J. et al. A framework for evaluating the performance of SMLM cluster analysis algorithms. Nature methods 20, 259–267, doi: 10.1038/s41592-022-01750-6 (2023).

15 Levet, F. et al. SR-Tesseler: a method to segment and quantify localization-based super-resolution microscopy data. Nat Methods 12, 1065–1071, doi:10.1038/nmeth.3579 (2015).

16 Williamson, D. J. et al. Machine learning for cluster analysis of localization microscopy data. Nat Commun 11, 1493, doi:10.1038/s41467-020-15293-x (2020).

17 Bender, S. W. B., Dreisler, M. W., Zhang, M., Kæstel-Hansen, J. & Hatzakis, N. S. SEMORE: SEgmentation and MORphological fingErprinting by machine learning automates super-resolution data analysis. Nature Communications 15, 1763, doi:10.1038/s41467-024-46106-0 (2024).

18 Saavedra, L. A., Mosqueira, A. & Barrantes, F. J. A supervised graph-based deep learning algorithm to detect and quantify clustered particles. Nanoscale 16, 15308–15318, doi:10.1039/D4NR01944J (2024).

19 Chen, T., Kornblith, S., Norouzi, M. & Hinton, G. A Simple Framework for Contrastive Learning of Visual Representations. Proceedings of the 37th International Conference on Machine Learning. 119, 1597–1607 (2020).

20 Becht, E. et al. Dimensionality reduction for visualizing single-cell data using UMAP. Nature Biotechnology 37, 38–44, doi:10.1038/nbt.4314 (2019).

21 McInnes, L., Healy, J. & Melville, J. UMAP: Uniform Manifold Approximation and Projection for Dimension Reduction. arXiv arXiv:1802.03426 (2018).

22 Shirgill, S. et al. Nano-org, a functional resource for single-molecule localisation microscopy data. Nature Communications 16, 8674, doi:10.1038/s41467-025-63674-x (2025).

23. Perlin, K. Improving noise. ACM Transactions on Graphics (TOG) 21, 681–682, doi:10.1145/566654.566636 (2002).

24 Perlin, K. An Image Synthesizer. Proceedings of the 12th Annual Conference on Computer Graphics and Interactive Techniques (SIGGRAPH 1985) 287–296 (Association for Computing Machinery, 1985).

25 De Brabander, M., De May, J., Joniau, M. & Geuens, G. Ultrastructural immunocytochemical distribution of tubulin in cultured cells treated with microtubule inhibitors. Cell Biol Int Rep 1, 177–183, doi:10.1016/0309-1651(77)90038-8 (1977).

26 von Diezmann, L., Shechtman, Y. & Moerner, W. E. Three-Dimensional Localization of Single Molecules for Super-Resolution Imaging and Single-Particle Tracking. Chem Rev 117, 7244–7275, doi:10.1021/acs.chemrev.6b00629 (2017).

27 Misra, D. Mish: A Self Regularized Non-Monotonic Activation Function. Proceedings of the British Machine Vision Conference (BMVC 2020), doi:10.5244/C.34.191 (2020).

28 Loshchilov, I. & Hutter, F. Decoupled Weight Decay Regularization. International Conference on Learning Representations (ICLR) (2019).

29 Hubert, L. & Arabie, P. Comparing partitions. Journal of classification 2, 193–218, doi: 10.1007/BF01908075 (1985).

30. Ester, M., Kriegel, H.-P., Sander, J. & Xu, X. A Density-Based Algorithm for Discovering Clusters in Large Spatial Databases with Noise. Proceedings of the Second International Conference on Knowledge Discovery and Data Mining (KDD 1996) 226–231 (AAAI Press) (1996).

31. Ripley, B. D. Modelling Spatial Patterns. Journal of the Royal Statistical Society: Series B 39, 172–192, doi: 10.1111/j.2517-6161.1977.tb01615.x (1977).

32 Paszke, A. et al. PyTorch: An Imperative Style, High-Performance Deep Learning Library. Advances in Neural Information Processing Systems, 32, 8024–8035 (Curran Associates, Inc.) (2019).

33 Pedregosa, F. et al. Scikit-learn: Machine Learning in Python. Journal of Machine Learning Research 12, 2825–2830 (2011).

34 Ries, J. SMAP: a modular super-resolution microscopy analysis platform for SMLM data. Nat Methods 17, 870–872, doi:10.1038/s41592-020-0938-1 (2020).

35 Schnitzbauer, J., Strauss, M. T., Schlichthaerle, T., Schueder, F. & Jungmann, R. Super-resolution microscopy with DNA-PAINT. Nature Protocols 12, 1198–1228, doi:10.1038/nprot.2017.024 (2017).

36 Edelstein, A. D. et al. Advanced methods of microscope control using μManager software. Journal of Biological Methods 1, e10, doi:10.14440/jbm.2014.36 (2014).

