## Supplementary Information for "Nanoscale spatial-omics via contrastive embedding of single-molecule localisation data"

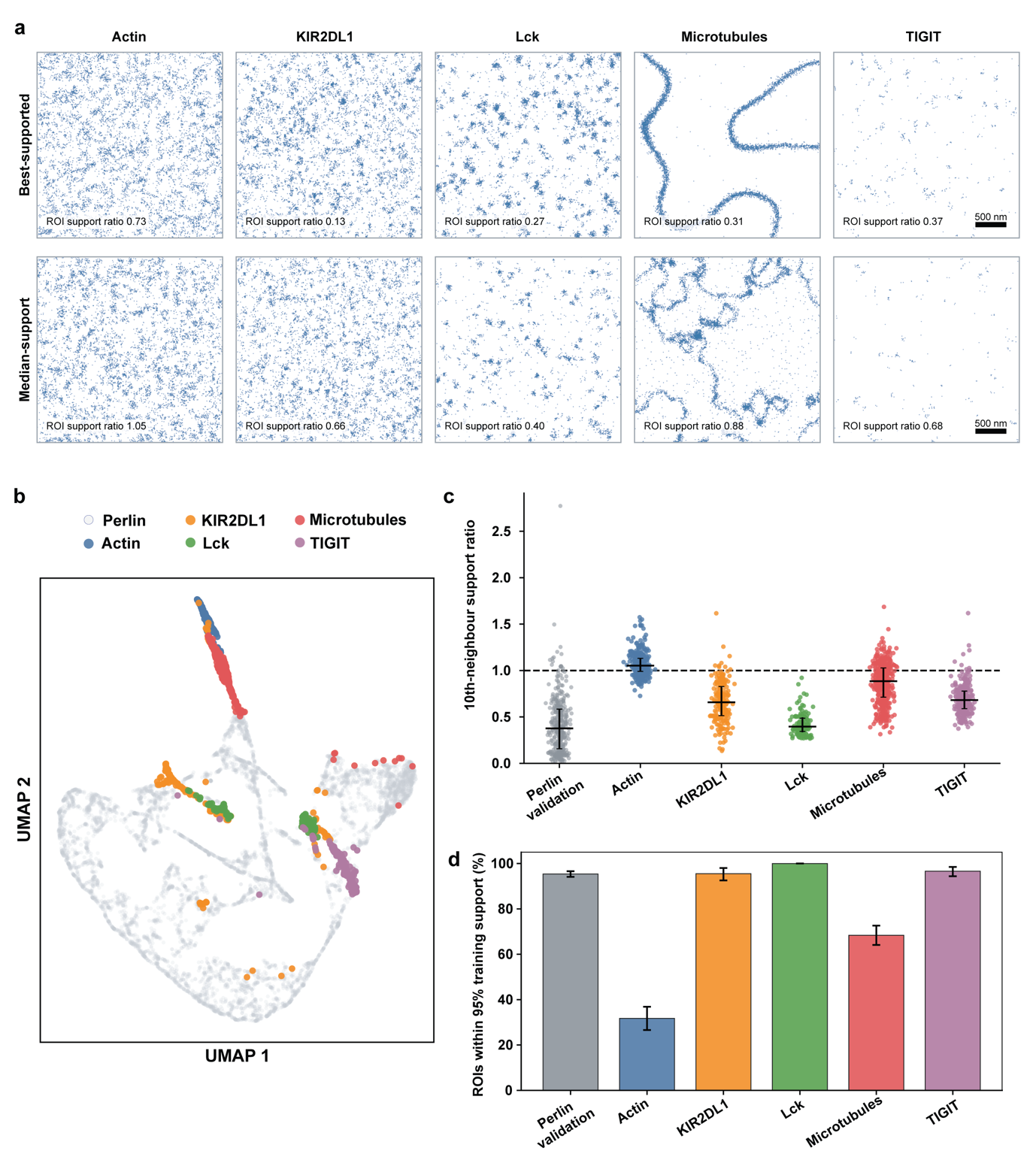

***Supplementary Figure S1 | Coverage of experimental SMLM point patterns by the synthetic Perlin training distribution.*** **a)** Representative Perlin training patterns selected according to their proximity to experimental protein ROIs in the learned 128-dimensional representation. For each protein (actin, KIR2DL1, Lck, microtubules and TIGIT), two training examples are shown: one selected as the closest available Perlin representation to the experimental ROI with the lowest training-support ratio (best-supported ROI; top), and one corresponding to an experimental ROI with approximately median training support (bottom). The annotated support ratios refer to the experimental ROIs used for selection. Patterns were reconstructed for visualisation from stored 25-nm localisation histograms, with positions within each bin randomly assigned. Each field represents a 3x3 µm ROI; scale bars, 500 nm. **b)** Joint UMAP projection of 8,000 Perlin training representations (light grey) and experimental SMLM ROIs from the five protein datasets (coloured). UMAP was used for visualisation only. **c)** Quantitative assessment of training-distribution support using cosine distances in the L2-normalised 128-dimensional representation. For each experimental ROI or independently generated Perlin validation sample, the distance to its tenth-nearest training representation was divided by the 95th percentile of tenth-nearest-neighbour distances among training representations, calculated excluding self-matches. The dashed line indicates a support ratio of 1, corresponding to the training-derived threshold. Values below 1 indicate representation-space neighbourhood distances within this threshold. Points represent individual ROIs or validation samples; black horizontal and vertical lines indicate the median and interquartile range, respectively. For visual clarity, 350 of the 1,000 independent Perlin validation samples are displayed, but summary statistics were calculated using the full validation dataset. **d)** Percentage of experimental ROIs and held-out Perlin validation samples satisfying the training-support criterion. Bars indicate the proportion within support, with error bars representing bootstrap 95% confidence intervals. The independent Perlin validation set (light grey; n = 1,000) provides an in-distribution reference, with 95.4% of samples falling within the training-derived threshold. Together, these analyses assess coverage within the learned representation rather than establishing exhaustive sampling of possible nanoscale spatial organisations. The experimental dataset comprised 1,352 ROIs: microtubules (n = 446), actin (n = 312), KIR2DL1 (n = 202), TIGIT (n = 267) and Lck (n = 125).

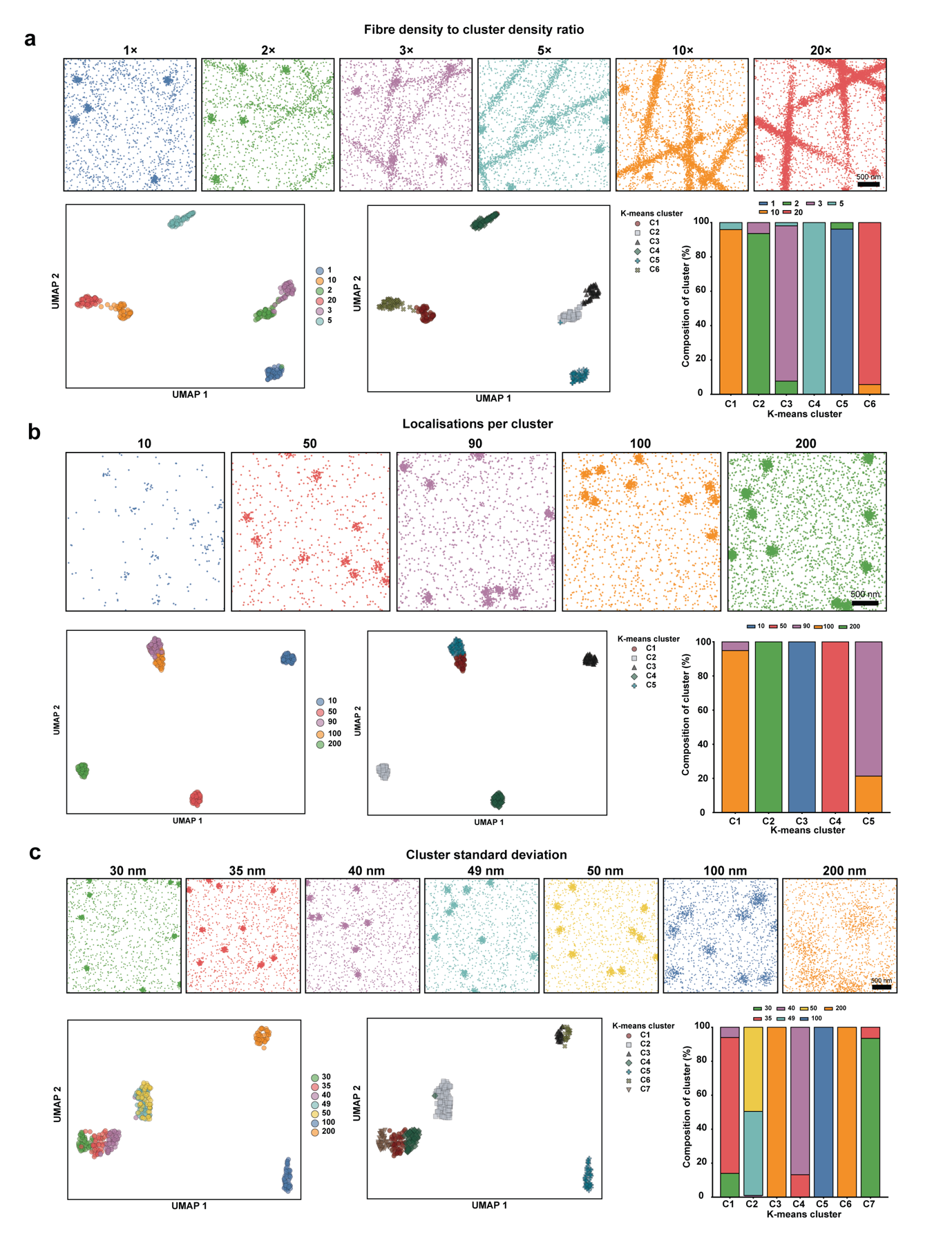

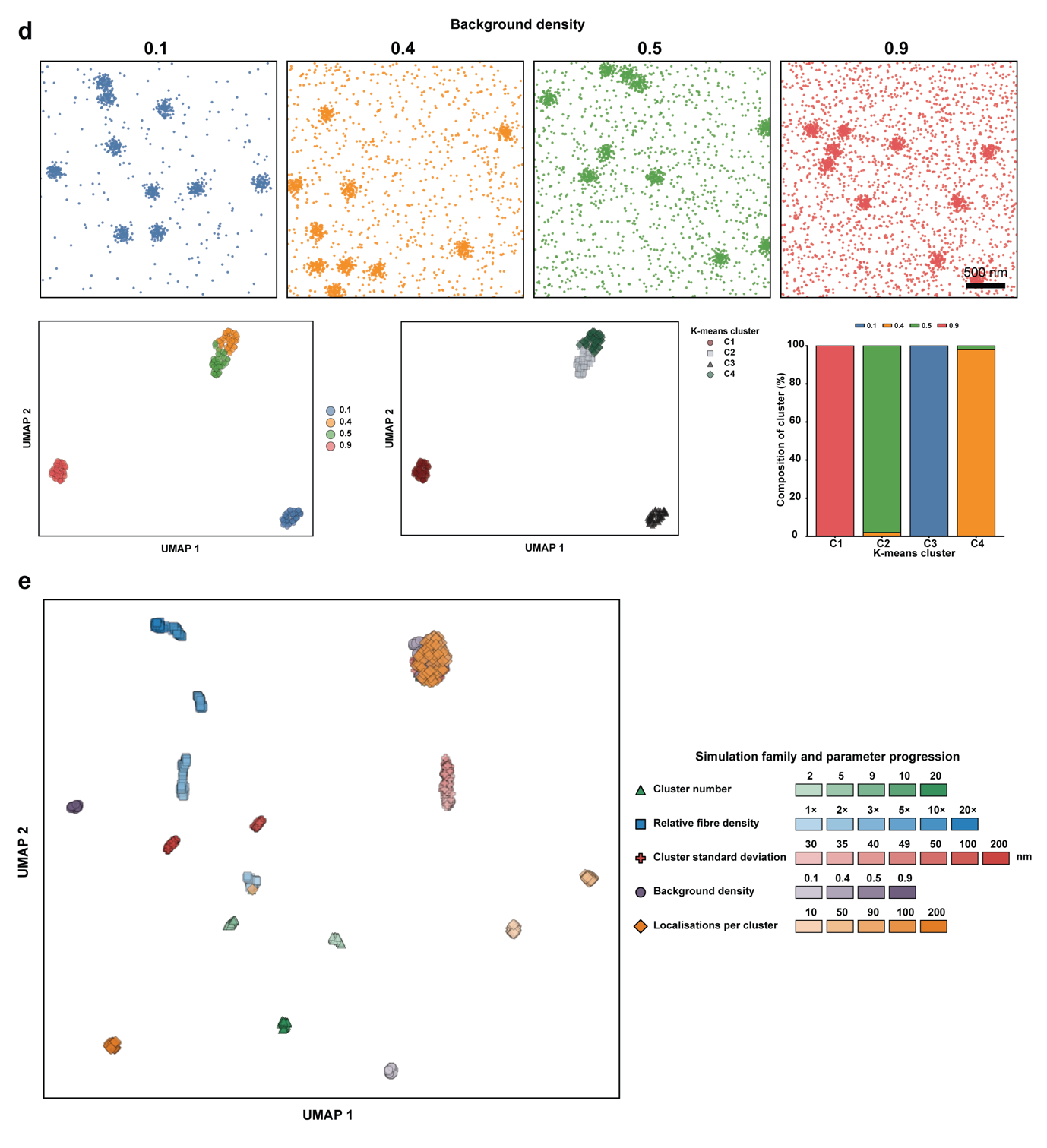

***Supplementary Figure S2: Response of the learned representation to controlled changes in simulated nanoscale organisation***. Simulated 3x3 µm ROIs were generated while systematically varying individual structural parameters. For each parameter sweep, representative point patterns are shown above a condition-coloured UMAP projection, a K-means-coloured UMAP projection and the percentage condition composition of each K-means cluster. K-means was performed in the full 128-dimensional representation with k equal to the number of simulated conditions; UMAP was used for visualisation only. **a)** Relative fibre density, defined as the number of localisations associated with fibre-like structures relative to Gaussian clusters, varied from 1x to 20x. **b)** Number of localisations per Gaussian cluster, varied from 10 to 200. **c)** Gaussian-cluster standard deviation, varied from 30 to 200 nm. **d)** Background-density parameter, varied from 0.1 to 0.9. Scale bars in representative simulations, 500 nm. **e)** Joint UMAP projection of all simulation families embedded together. Marker shape denotes simulation family and colour shade denotes the parameter level within each family, as indicated by the accompanying swatches. This joint projection illustrates relationships among simulation families in a common learned representation. Each simulated condition comprised 50 independently generated ROIs. The individual parameter sweeps included 300 ROIs for **a)** relative fibre density, 250 for **b)** localisations per cluster, 350 for **c)** Gaussian-cluster standard deviation and 200 for **d)** background variation. The joint analysis e) included these four simulation families together with the Gaussian-cluster-number simulation from **Figure 2**, giving five simulation families and 1,350 ROIs in total.

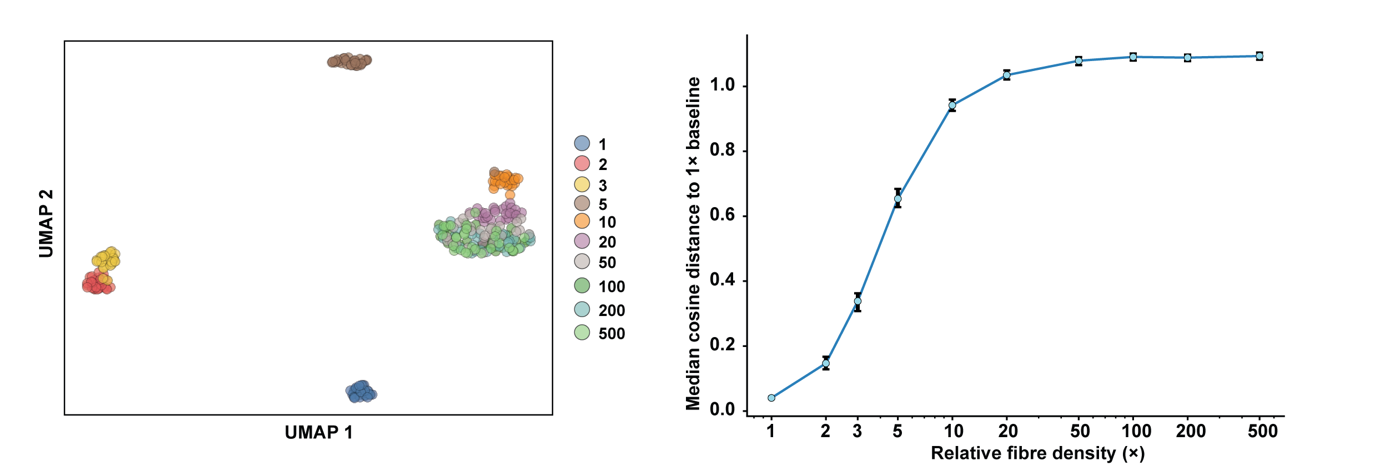

***Supplementary Figure S3: Saturation of the learned response to increasing relative fibre density***. Independent simulated 3x3 µm ROIs containing Gaussian clusters and fibre-like structures were generated at relative fibre-density multipliers of 1x, 2x, 3x, 5x, 10x, 20x, 50x, 100x, 200x and 500x (n = 30 ROIs per condition, total = 300 ROIs). Left, UMAP visualisation of the resulting 128-dimensional representations, coloured by relative fibre density. Right, median cosine distance between each condition and the 1x reference distribution in the normalised 128-dimensional representation; error bars show bootstrap 95% confidence intervals. The response increases strongly at low-to-intermediate fibre densities and approaches a plateau at high fibre densities.

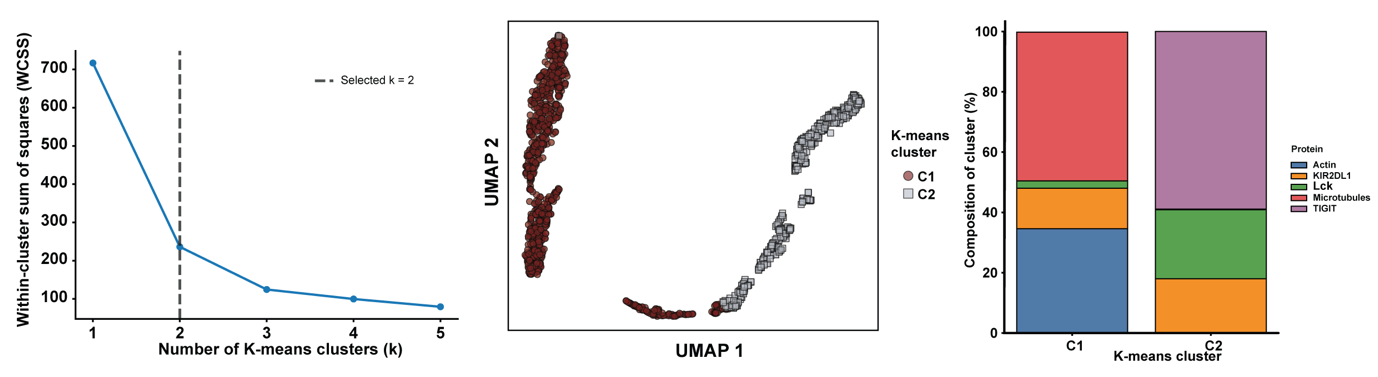

***Supplementary Figure S4: Unconstrained estimation of cluster number in the experimental protein dataset***. Left, within-cluster sum of squares (WCSS) across candidate values of k for the five-protein dataset shown in **Figure 3**; the elbow criterion selected k = 2. Centre, UMAP visualisation coloured by the resulting two K-means clusters. Right, percentage protein composition of each cluster. K-means clustering and calculation of WCSS were performed in the full 128-dimensional representation; UMAP was used only for visualisation. The analysis included the same 1,352 experimental ROIs used in **Figure 3**.

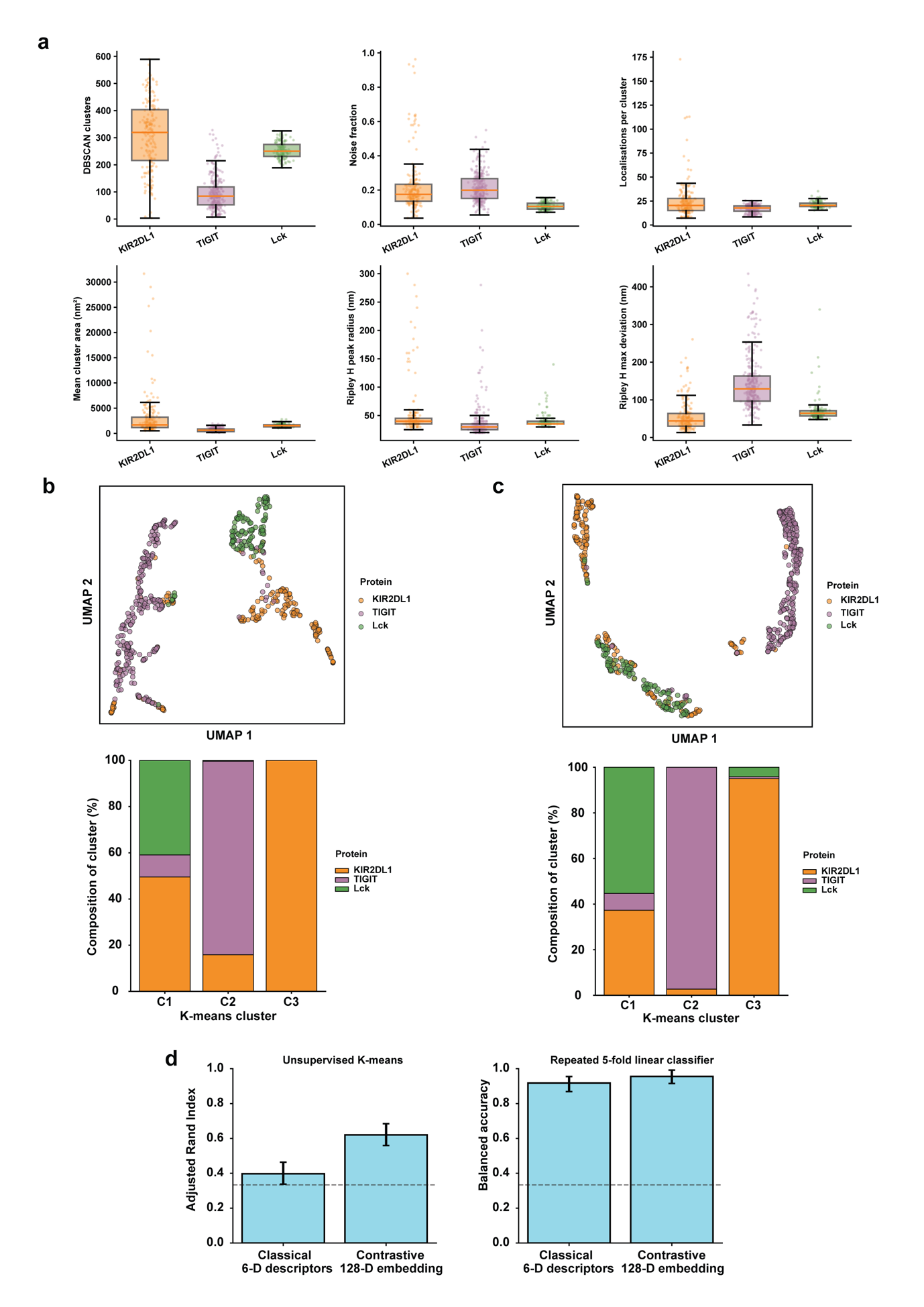

***Supplementary Figure S5: Head-to-head comparison of predefined spatial descriptors and the learned contrastive representation***. Analysis was performed on the non-fibrous KIR2DL1, TIGIT and Lck datasets using identical ROIs. **a)** Six conventional spatial descriptors calculated independently for each ROI: DBSCAN cluster number, noise fraction, mean localisations per cluster and mean cluster area, together with Ripley-derived peak radius and maximum deviation. DBSCAN was performed using ε = 25 nm and minimum points = 5. **b)** UMAP visualisation of the standardised six-dimensional classical-descriptor representation (top) and percentage condition composition after K-means clustering with k = 3 in the full six-dimensional descriptor space (bottom). **c)** Corresponding UMAP visualisation and K-means composition obtained from the 128-dimensional contrastive representation of the same ROIs; K-means was performed with k = 3 in the full representation. **d)** Head-to-head assessment of condition discriminability. Left, Adjusted Rand Index (ARI) for unsupervised K-means clustering in the classical six-dimensional and contrastive 128-dimensional representations; error bars show bootstrap 95% confidence intervals. Right, balanced accuracy from a linear logistic-regression classifier evaluated by repeated stratified five-fold cross-validation; error bars show the 2.5th–97.5th percentile range of cross-validation scores. The analysis comprised 594 experimental ROIs: KIR2DL1 (n = 202), TIGIT (n = 267) and Lck (n = 125).

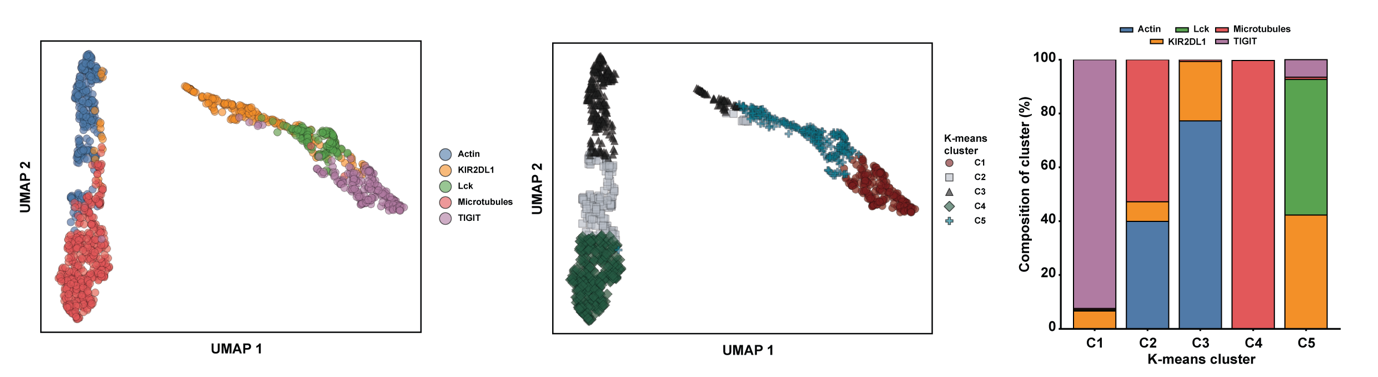

***Supplementary Figure S6: Protein organisation remains distinguishable after localisation-density normalisation.*** Experimental ROIs for microtubules, actin, KIR2DL1, TIGIT and Lck were randomly subsampled to a fixed density of 100 localisations µm⁻² before encoding. Left, UMAP projection coloured by protein identity. Centre, the same ROIs coloured by K-means cluster assignment. Right, percentage protein composition of each K-means cluster. K-means clustering was performed in the full 128-dimensional representation with k = 5. Following exclusion of ROIs with fewer than 900 localisations, the density-normalised dataset comprised 1,296 ROIs: microtubules (n = 446), actin (n = 312), KIR2DL1 (n = 200), TIGIT (n = 213) and Lck (n = 125).

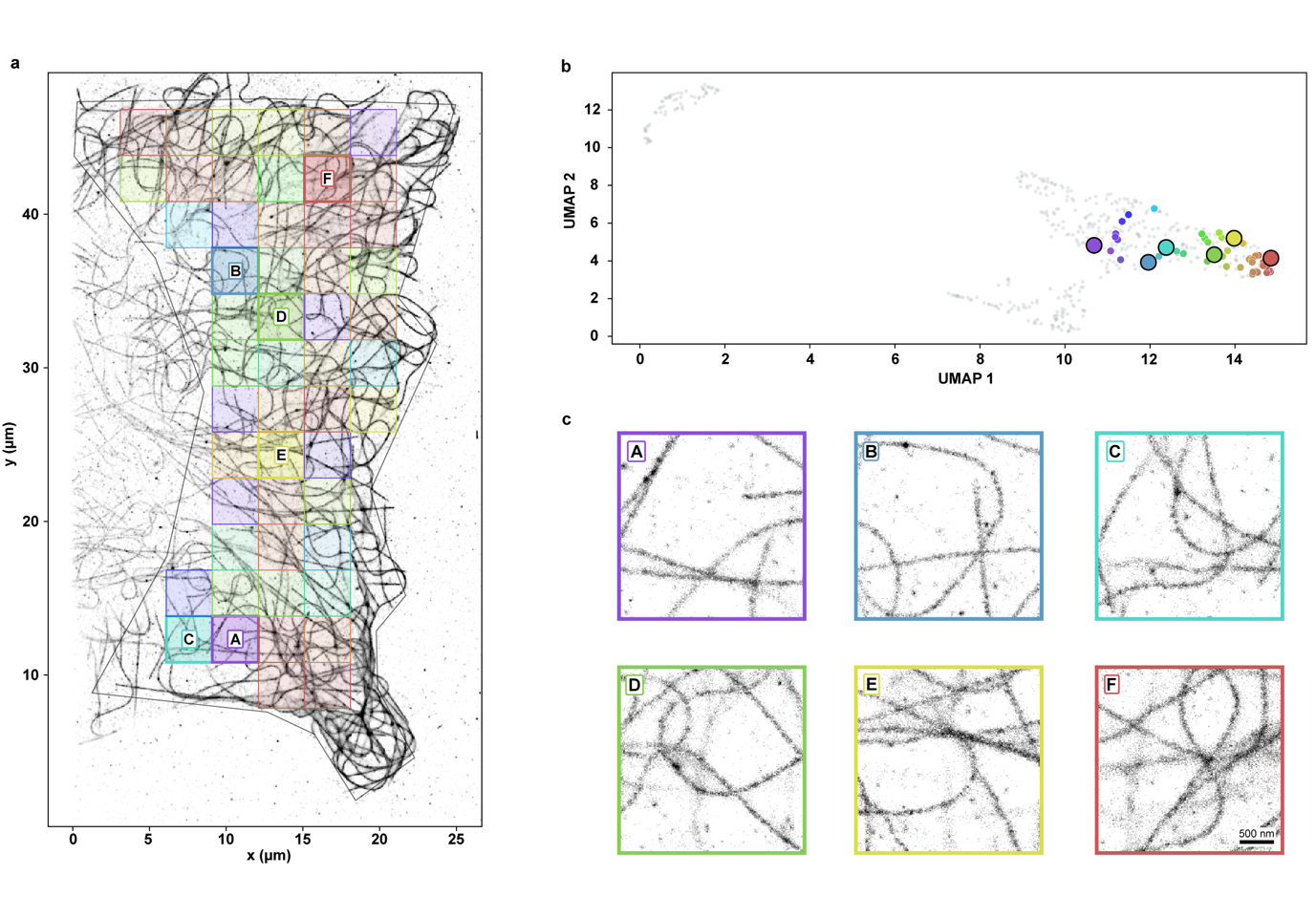

***Supplementary Figure S7: Mapping microtubule organisation within a single cell to the learned representation***. **a)** SMLM reconstruction of microtubules in a representative AF647-labelled COS-7 cell, overlaid with the 3x3 µm ROI grid used for contrastive analysis (n = 52 ROIs). Individual ROIs are coloured according to their positions in the two-dimensional UMAP projection, with six representative ROIs (**A–F**) highlighted. **b)** UMAP visualisation of the 128-dimensional representations generated using the pretrained Pool8 encoder. Grey points indicate the broader experimental reference embedding from **Figure 4**, while coloured points correspond to the individual ROIs from the cell shown in **a)**. Enlarged markers identify the six representative ROIs (**A–F**) selected across the occupied projection region. **c)** Enlarged SMLM reconstructions of the six selected ROIs, illustrating variation in local microtubule organisation within a single cell. Matching colours link each ROI to its position in the **a)** full-cell reconstruction and the **b)** UMAP projection. Scale bar, 500 nm.

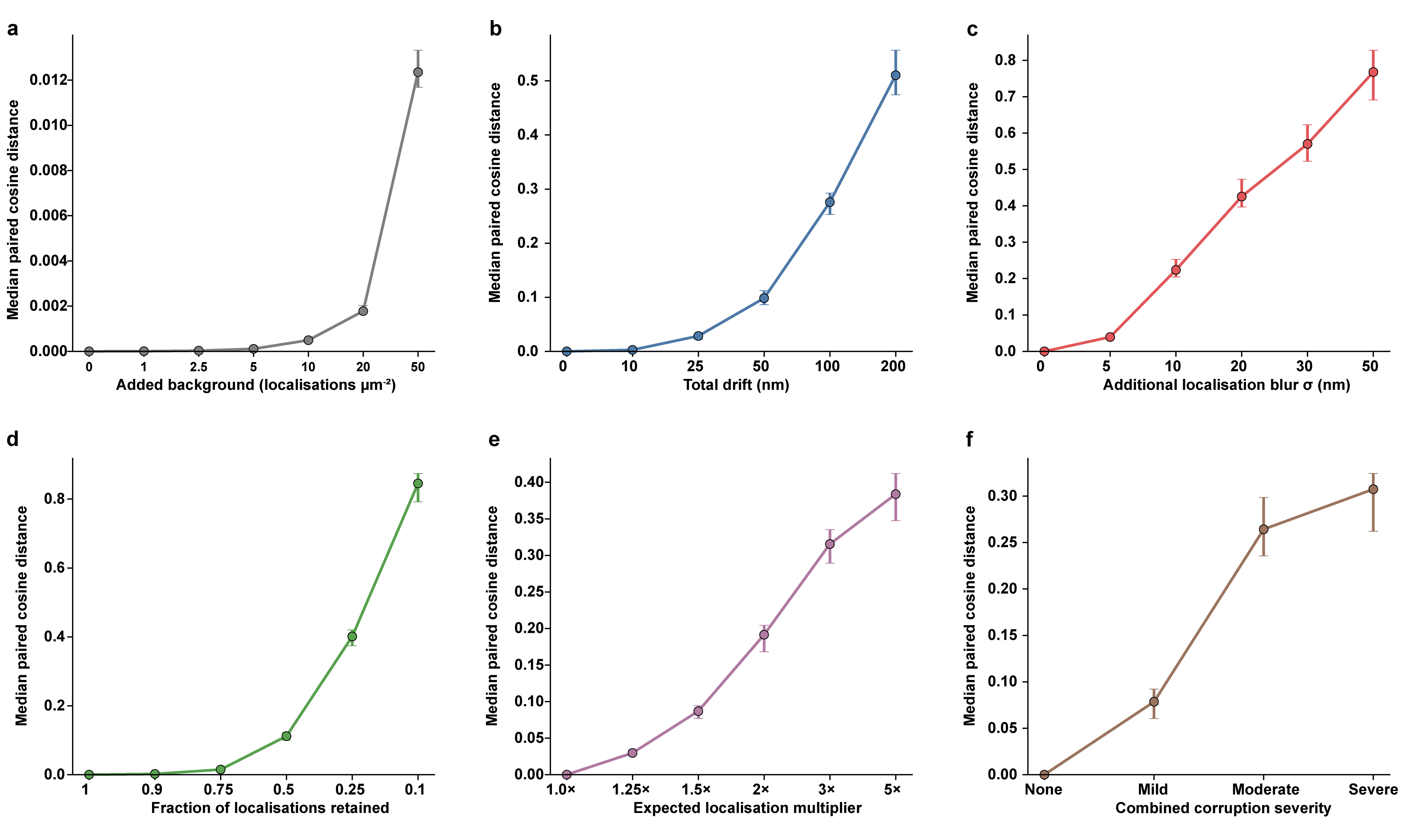

***Supplementary Figure S8: Sensitivity of the learned representation to graded technical perturbations of SMLM data***. A set of 125 experimental Lck ROIs was subjected to systematic in silico corruption, with each perturbed ROI compared directly with the representation of the same unmodified ROI. Curves show the median paired cosine distance in the normalised 128-dimensional representation; error bars indicate bootstrap 95% confidence intervals. **a)** Addition of uniform background localisations at 1–50 localisations µm⁻². **b)** Linear drift with total first-to-last displacement of 10–200 nm. **c)** Additional Gaussian localisation blur with σ = 5–50 nm per coordinate axis. **d)** Random thinning retaining 90%, 75%, 50%, 25% or 10% of the original localisations. **e)** Simulated blinking/overcounting producing 1.25x, 1.5x, 2x, 3x or 5x the original number of localisations, with repeated localisations jittered by σ = 10 nm. **f)** Combined multi-artifact stress tests. The mild profile retained 85% of localisations and combined 5 nm blur, 25 nm drift, 1.25x overcounting and 2.5 added background localisations µm⁻²; the moderate profile retained 65% and combined 10 nm blur, 50 nm drift, 1.5x overcounting and 5 background localisations µm⁻²; the severe profile retained 40% and combined 20 nm blur, 100 nm drift, 2x overcounting and 10 background localisations µm⁻². For all perturbation families, the median within-ROI Spearman correlation between perturbation severity and representation displacement was ρ = 1.00.

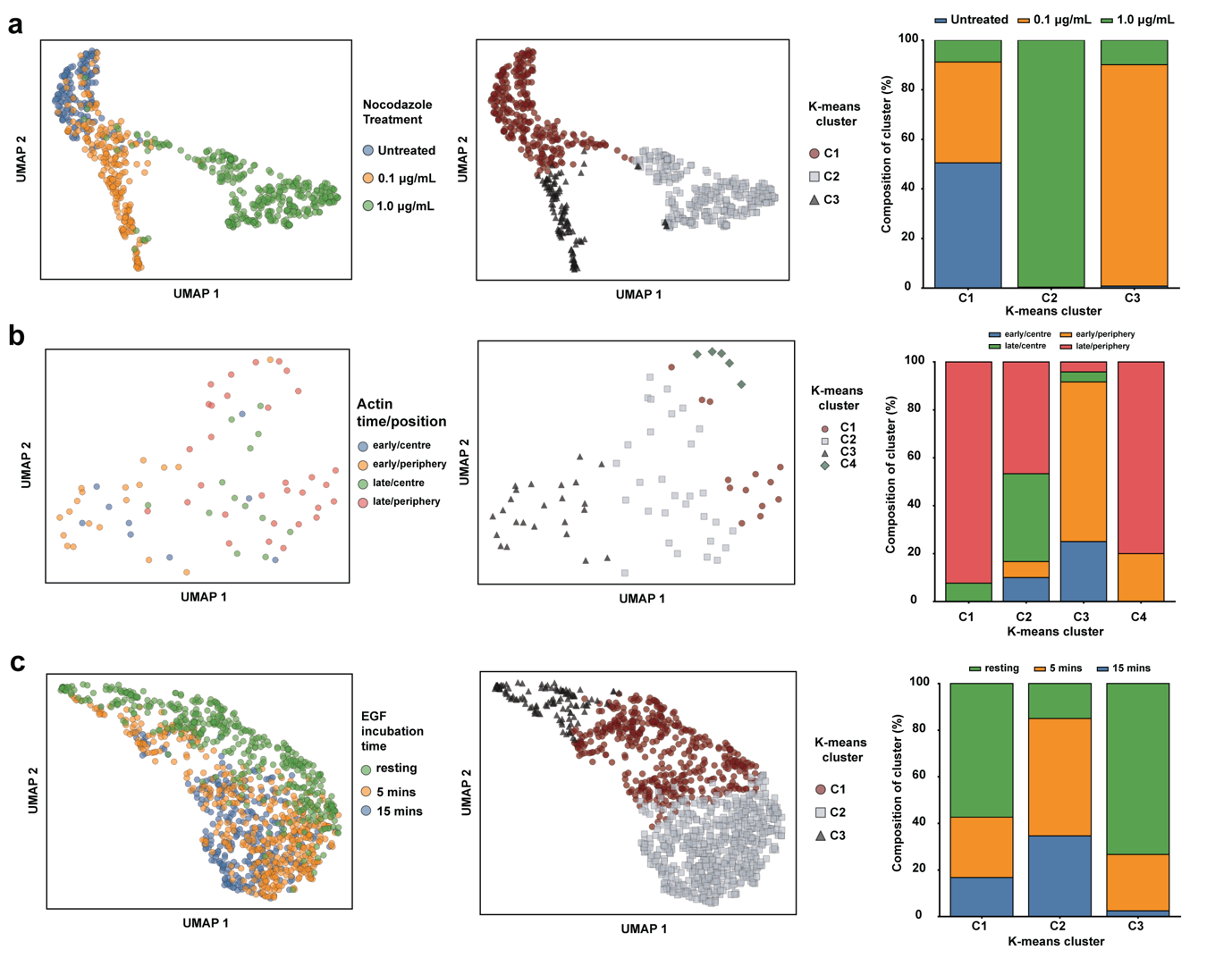

***Supplementary Figure S9: Additional biological perturbation analyses and localisation-density control***. **a)** Nocodazole-treated microtubule ROIs were randomly subsampled to 100 localisations µm⁻² before analysis to assess whether the dose-dependent separation observed in Figure 5 persisted after localisation-density normalisation. The analysis retained all 709 ROIs: untreated control (n = 150), 0.1 µg mL⁻¹ nocodazole (n = 229) and 1.0 µg mL⁻¹ nocodazole (n = 330). **b)** Actin SMLM ROIs from Jurkat E6.1 immunological synapses stratified by synapse stage and spatial region, comprising early centre (n = 9), early periphery (n = 19), late centre (n = 13) and late periphery (n = 31), for a total of 72 ROIs. **c)** EGFR SMLM ROIs from HeLa cells under resting conditions (n = 428) or following EGF stimulation for 5 min (n = 444) or 15 min (n = 284), for a total of 1,156 ROIs. For each analysis, panels show a condition-coloured UMAP projection, the corresponding K-means-cluster projection and percentage condition composition of each K-means cluster. K-means clustering was performed in the full 128-dimensional representation with k equal to the number of experimental conditions.
